# Ovothiol A mediates singlet oxygen resistance and acclimation in Chlamydomonas

**DOI:** 10.64898/2026.02.12.702910

**Authors:** Yuliia Lihanova, Félix de Carpentier, Gladwin Suryatin Alim, Elisabeth Hommel, Matthias Hirth, Gabriella Benko, Sanoja C. Sridevan, Raimund Nagel, Matthias Gilbert, Christian Hertweck, Arthur R. Grossman, Florian P. Seebeck, Krishna K. Niyogi, Setsuko Wakao, Severin Sasso

## Abstract

Light is essential for photosynthetic organisms, but excess light can generate toxic levels of reactive oxygen species (ROS). To neutralize these ROS, plants and algae produce a variety of antioxidants like carotenoids, tocopherols, and glutathione. However, the role of alternative ROS scavengers, such as ovothiols, has not been studied in the context of oxidative stress in photosynthetic organisms. Here, we report that many algal groups have the potential for the biosynthesis of ovothiols, a group of thiohistidines. We discovered that the model green microalga *Chlamydomonas reinhardtii* produces millimolar concentrations of ovothiol A, whose biosynthesis is mediated by the ovothiol synthase OVOA1. Using CRISPR-generated *ovoa1* knockout mutants, we found that ovothiol production is essential for resistance and acclimation to singlet oxygen, a prominent ROS in photosynthetic organisms. Finally, we demonstrated that *OVOA1* expression is activated by singlet oxygen and light signaling pathways in which we identified the major regulatory factors. Overall, our results show that ovothiol A is a major, previously overlooked antioxidant in Chlamydomonas. This work broadens our understanding of cellular mechanisms that combat the damaging effects of oxidative stress.

**Graphical abstract:** 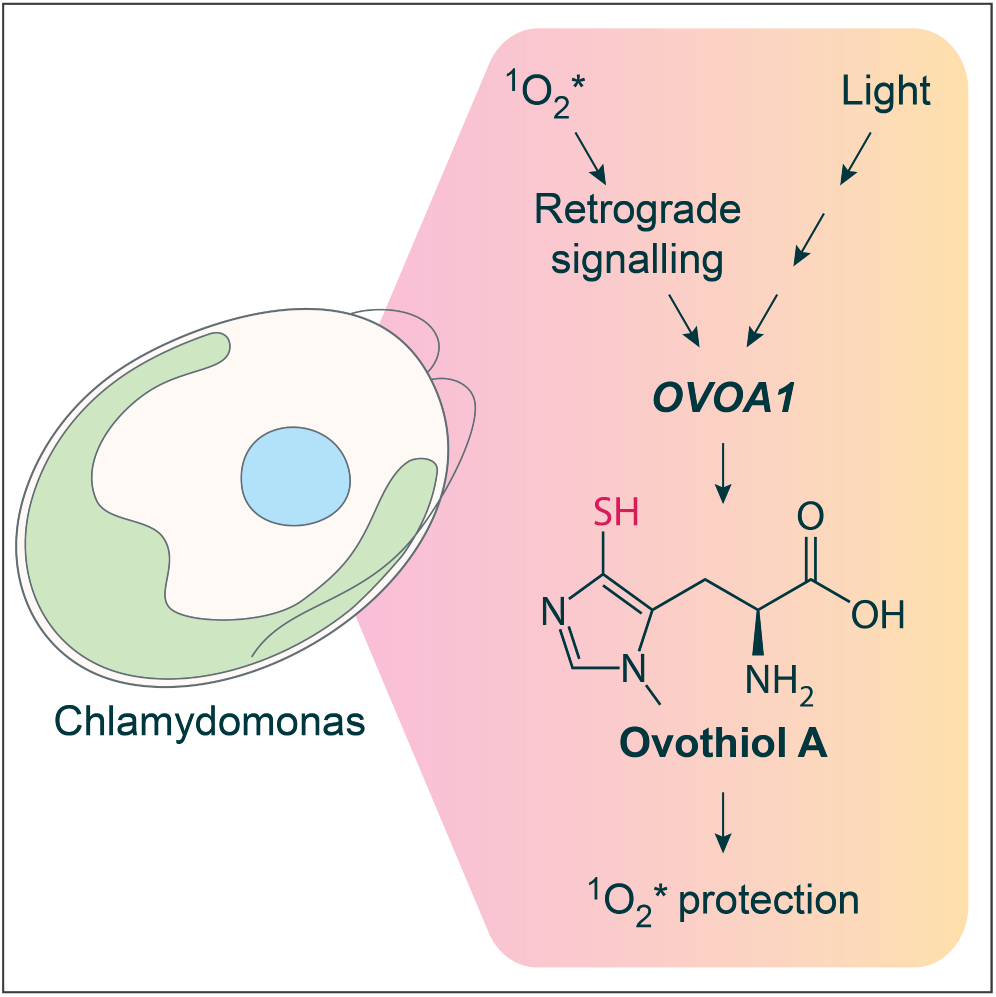

## Introduction

Plants and algae must overcome the challenge that using light as a source of energy in the presence of oxygen poses a constant risk of cell damage. Exposure of photosynthetic organisms to high light induces the production of reactive oxygen species (ROS) when the photon flux saturates the chloroplast electron transport chain. Superoxide is generated at the acceptor side of photosystem I and converted to hydrogen peroxide by superoxide dismutase. If it is not promptly converted to water, hydrogen peroxide could lead to the generation of hydroxyl radicals through non-enzymatic reactions (reviewed in Mittler et al. 2022). In contrast, singlet oxygen (^1^O_2_*) is generated predominantly in the reaction center of photosystem II when the electron transport chain is over-reduced and the yield of triplet excited chlorophyll molecules increases (reviewed in Krieger-Liszkay et al. 2008). ^1^O_2_* can also be induced in the dark by other stress factors or originate from the environment (Prasad et al. 2016; Larson and Marley 1999). The accumulation of ROS ultimately results in the damage of cellular proteins, membranes, and nucleic acids (Triantaphylidès et al. 2008; Barciszewski et al. 1999; Babbar et al. 2021). In response to oxidative stress plants and microalgae produce a variety of antioxidants (reviewed in Wakao and Niyogi 2021; Mittler et al. 2022). ^1^O_2_* can be directly scavenged by tocopherols (vitamin E), plastoquinol, and carotenoids (Krieger-Liszkay et al. 2008; Li et al. 2012; Nowicka and Kruk 2012). In addition, other mechanisms have evolved by which the unicellular model alga *Chlamydomonas reinhardtii* (hereafter referred to as “Chlamydomonas”) acclimates to ^1^O_2_*. Exposure of Chlamydomonas to a low dose of ^1^O_2_* elicits expression of a subset of genes which increases the resistance of the alga to subsequently applied higher doses (Ledford et al. 2007). In this case, ^1^O_2_* initially acts in retrograde signaling through the regulatory protein SAK1 (SINGLET OXYGEN ACCLIMATION KNOCKED-OUT1; Wakao et al. 2014). On the other hand, hydrogen peroxide is mainly detoxified by the action of multiple antioxidants including glutathione and ascorbate together with glutathione and ascorbate peroxidases (reviewed in Wakao and Niyogi 2021). Whereas the antioxidant glutathione has been extensively studied, the roles of other thiol antioxidants, including the ovothiols, have been largely overlooked in photosynthetic eukaryotes (Castellano and Seebeck 2018).

Ovothiols are small thiohistidine antioxidants with variable methylation levels of their α-amino group (Figure 1A): ovothiol A is not methylated whereas ovothiols B and C are mono- and dimethylated, respectively. The initial step in ovothiol A biosynthesis (Figure 1B) is mediated by the sulfoxide synthase domain of ovothiol synthase (OvoA), which conjugates a cysteine and a histidine to form a 5-histidylcysteine sulfoxide (Braunshausen and Seebeck 2011). The latter is cleaved by the β-lyase OvoB to form 5-thiohistidine, whose imidazole ring is finally methylated by the OvoA methyltransferase domain, yielding ovothiol A (Naowarojna et al. 2018). The diversity of organisms producing ovothiols ranges from bacteria and trypanosomes to invertebrates and algae. In the green alga *Dunaliella salina*, ovothiol A disulfide may have a regulatory function as it inactivates the plastid ATP synthase complex *in vitro* (Selman-Reimer et al. 1991). In addition, ovothiols have been detected in diatoms and *Euglena gracilis*, but their function remains enigmatic (O’Neill et al. 2015; Milito et al. 2020a). The role of ovothiols as antioxidants has only been described in non-photosynthetic organisms (reviewed in Castellano and Seebeck 2018), including protecting against oxidative stress during the fertilization of sea urchin eggs (Turner et al. 1988). As for unicellular organisms, ovothiols protect trypanosomatids from oxidative stress during infection (Ariyanayagam and Fairlamb 2001). This protective action of ovothiols is attributed to two antioxidant properties. First, due to the very low *pKa* of their thiol group, ovothiols readily react with hydrogen peroxide under physiological conditions to form ovothiol disulfide, which is then spontaneously reduced by glutathione via thiol-disulfide exchanges (Turner et al. 1988; Osik et al. 2021). Second, ovothiols can directly scavenge radicals such as superoxide or hydroxyl radical, and also quench the triplet state of molecules like kynurenic acid (Holler and Hopkins 1990; Marjanovic et al. 1995; Osik et al. 2021). In contrast, it is not known if ovothiols are involved in detoxifying ^1^O_2_*. However, similar histidine-derived thiols, such as ergothioneine and 2-thiohistidine, as well as other imidazole-containing compounds can efficiently react with ^1^O_2_* (Dahl et al. 1988; Hartmann et al. 2023; Hartman et al. 1990). This ROS is particularly prevalent in the biology of photosynthetic organisms because it is generated as a consequence of photosynthetic activity and is the primary agent of photooxidative stress (Triantaphylidès et al. 2008).

**Figure 1.**
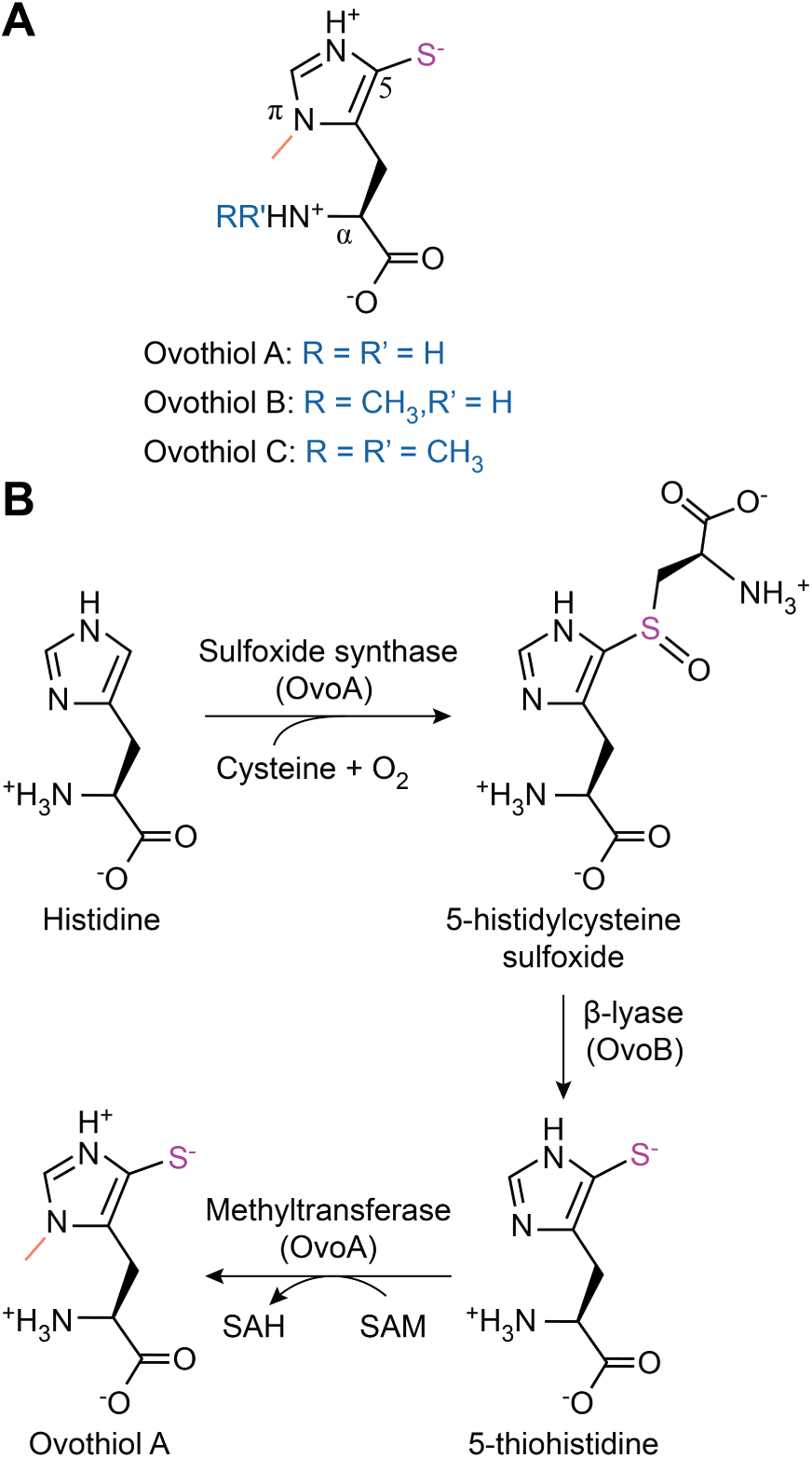
Chemical structure and biosynthesis of ovothiols. (A) Structure of the thiohistidines ovothiol A, B, and C. The α-amino group of ovothiol A is unmethylated, whereas ovothiols B and C are *N*-monomethylated and *N*,*N*-dimethylated, respectively (in blue). (B) Biosynthetic pathway of ovothiol A. First, the OvoA sulfoxide synthase domain catalyzes the conjugation of cysteine and histidine to form 5-histidylcysteine sulfoxide (Braunshausen and Seebeck 2011). Second, the sulfoxide conjugate is cleaved by the pyridoxal phosphate-dependent β-lyase OvoB resulting in 5-thiohistidine (sulfur atom in purple). Finally, the imidazole ring of 5-thiohistidine is methylated (in orange) by the OvoA methyltransferase domain, yielding the final product ovothiol A (Naowarojna et al. 2018). SAM, *S*-adenosyl methionine; SAH, *S*-adenosyl homocysteine.

In this study, we set out to elucidate the role of ovothiol in Chlamydomonas. In the first step of our investigation, we show that ovothiol A is produced in millimolar concentrations and is present in its reduced form. Using CRISPR-Cas9-generated mutants, we show that the ovothiol synthase OVOA1 participates in ovothiol A biosynthesis. Additionally, *OVOA1* was found to be essential for resistance and acclimation to ^1^O_2_*. Finally, we show that the expression of *OVOA1* is regulated by ^1^O_2_*-and light-induced signaling pathways. Together, these findings demonstrate that ovothiol A is a major antioxidant in Chlamydomonas, broadening our knowledge of oxidative stress responses in photosynthetic eukaryotes.

## Results

### Homologous genes encoding ovothiol synthase (OvoA) are widespread in algae

To investigate ovothiol production in photosynthetic eukaryotes, we focused first on identifying genes encoding the ovothiol biosynthetic enzyme OvoA (Figure 1B; Braunshausen and Seebeck 2011). In a maximum-likelihood phylogenetic tree of eukaryotic and bacterial sequences, OvoA homologs were found in the genomes of diverse algal groups including haptophytes, photosynthetic alveolates, ochrophytes, cryptophyceae, *Symbiodinium* species, diatoms, and Archaeplastida (Figure 2A). Most plant, animal, and stramenopile homologs are grouped together nested within the sequences from heterotrophic bacteria. This suggests that an ancient common ancestor of these eukaryotes acquired OvoA from heterotrophic bacteria through lateral gene transfer. This hypothesis is supported by the fact that archaeal, cyanobacterial, and alphaproteobacterial ancestors of eukaryotes, plastids, and mitochondria do not contain sequences similar to OvoA.

**Figure 2.**
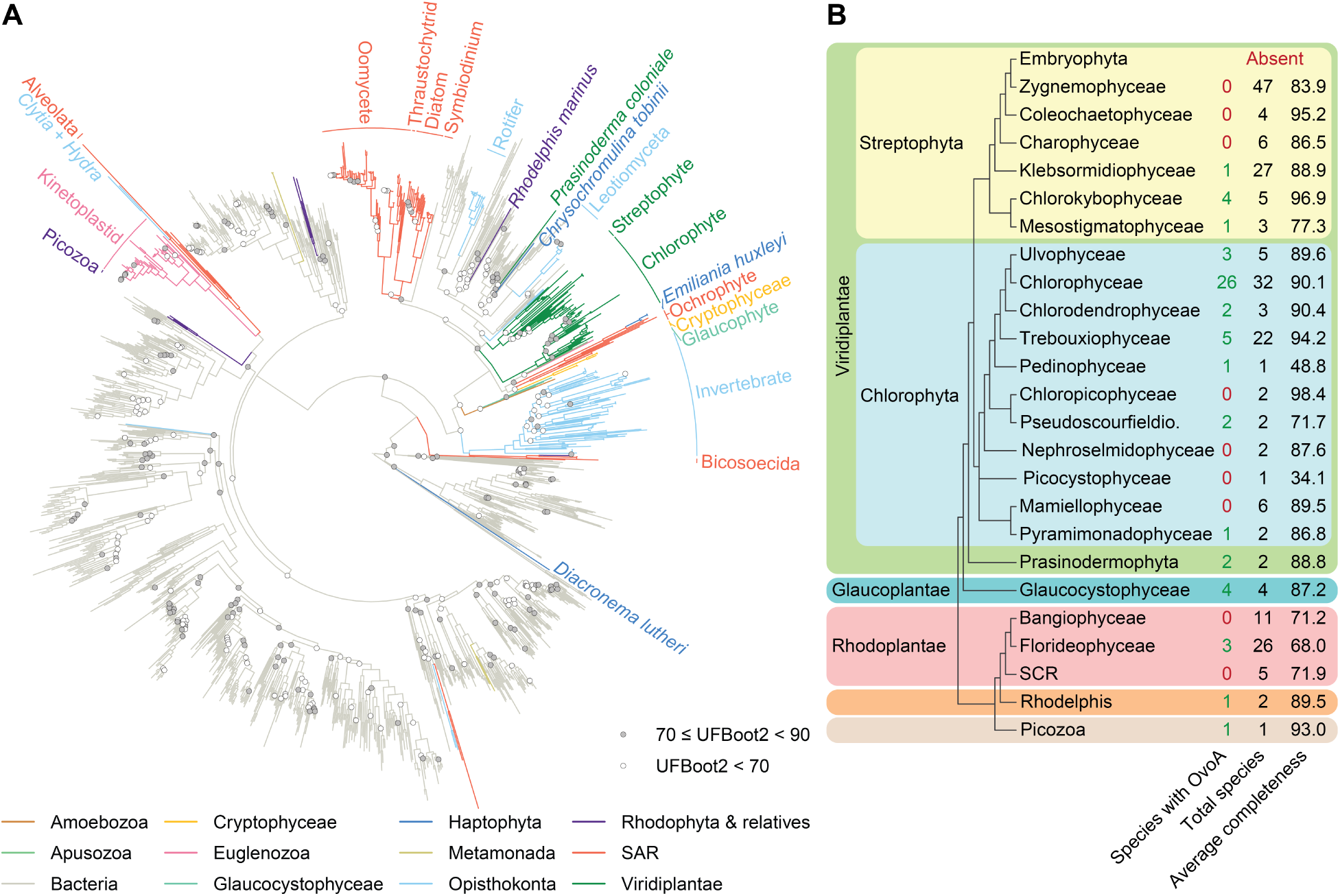
Occurrence and evolution of OvoA homologs in algae and plants. (A) Phylogeny of OvoA homologs with relevant eukaryotic clades highlighted. The maximum likelihood tree was generated with protein sequences using the Q.pfam+I+R10 model (Minh et al. 2021). The tree was rooted by minimal ancestor deviation (Tria et al. 2017) placing ergothioneine synthases as an outgroup (not shown in the figure). UFBoot2 branch support values above 90 are not shown. (B) Occurrence of OvoA homologs in Archaeplastida. Completeness for each species was estimated by BUSCO (Benchmarking Universal Single-Copy Homologs) using the OrthoDB 12 eukaryote database (Tegenfeldt et al. 2025) and averaged for all species of each taxon. SCR, Stylonematophyceae + Compsopogonophyceae + Rhodellophyceae; Pseudoscourfieldio, Pseudoscourfieldiophyceae.

We then analyzed the prevalence of genes encoding OvoA across Archaeplastida, which was estimated from a collection of genomes and transcriptomes (Figure 2B; Supplementary Table S1). Homologous genes were found in all major clades: chlorophytes, streptophytes, prasinodermophytes, glaucophytes, as well as rhodophytes and their non-photosynthetic relatives *Rhodelphis* and *Picozoa*. OvoA was present in basal streptophyte microalgae, but was lost in the monophyletic clade comprising Charophyceae, Coleochaetophyceae, Zygnematophyceae, and land plants. In Chlorophyta, OvoA was detected in most classes and especially enriched in Chlorophyceae, which includes *Dunaliella salina*, a known producer of ovothiol A disulfide (Selman-Reimer et al. 1991). Interestingly, it was present in the genome of Chlamydomonas, which prompted us to investigate ovothiols in this genetically tractable model green alga (Sasso et al. 2018).

### Chlamydomonas accumulates millimolar concentrations of reduced ovothiol A

Ovothiols exist in three different forms which differ in the methylation level of their α-amino group (Figure 1A). We sought to determine if Chlamydomonas produces any of those thiohistidines. In cells, thiols can be reduced, oxidized to disulfides, or bound to protein (Turner et al. 1988; Ariyanayagam and Fairlamb 2001; Zaffagnini et al. 2012). For easier identification, cellular thiols were fully reduced and derivatized with 4-bromomethyl-7-methoxycoumarin (BMC), which is suitable for metabolite identification by liquid chromatography-mass spectrometry (LC-MS) and identification and quantification by high-performance liquid chromatography (HPLC) coupled to absorbance measurements (Supplementary Figure S1A; Milito et al. 2020; Liao et al. 2025). Under constant low light conditions (50 µmol photons m^-2^ s^-1^), ovothiol A was detected with a clear peak of the expected ion *m/z*, whereas no ovothiol B or C was detected (Figures 3A and B). We then quantified the intracellular concentration of ovothiol A. As no ovothiol standard was available, 2-thiohistidine was used as the standard because of its similar structure (Supplementary Figure S2). Assuming a uniform intracellular distribution, ovothiol A accumulated at a concentration of 2.3 mM making it one of the most abundant thiol antioxidants in Chlamydomonas.

**Figure 3.**
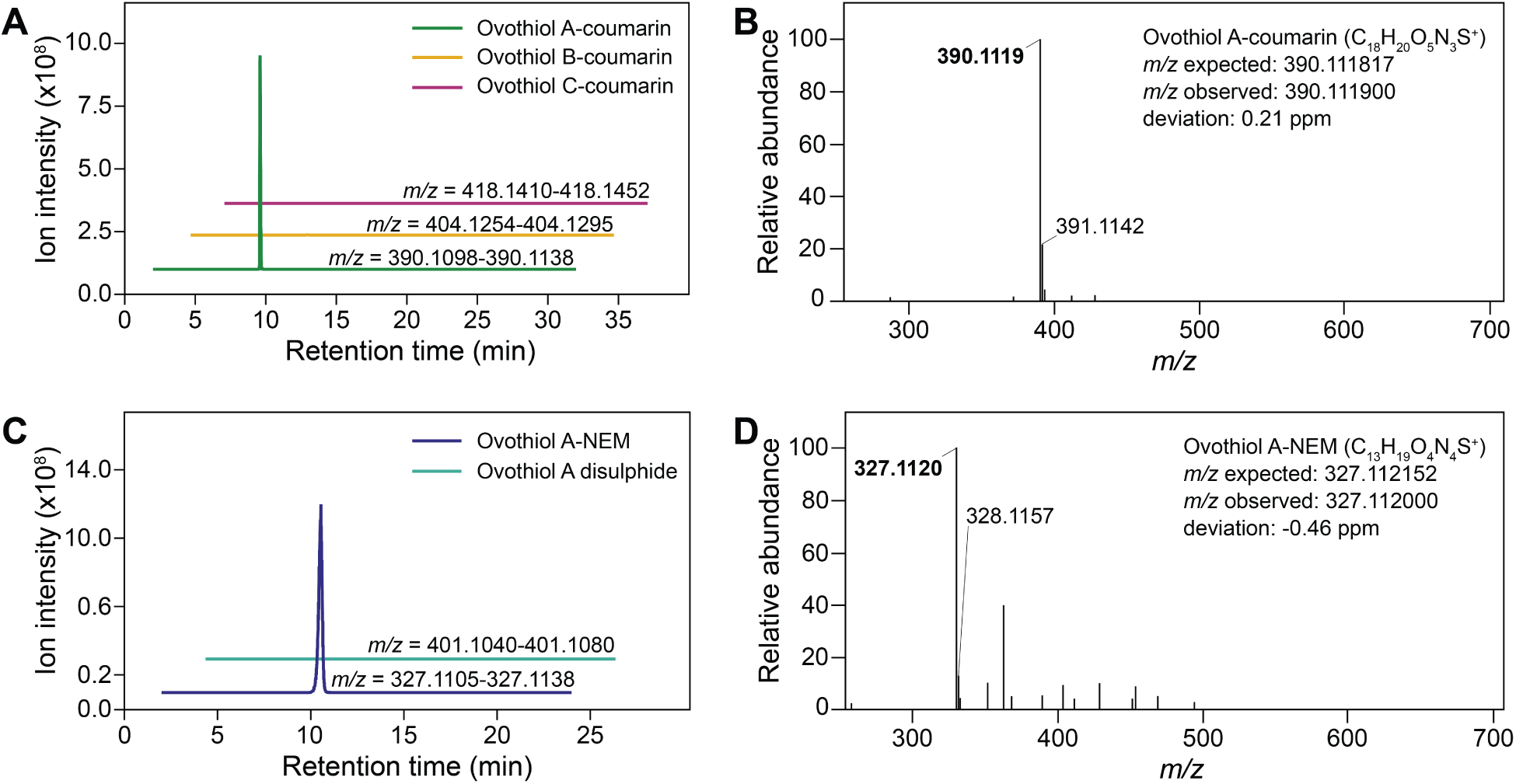
Chlamydomonas accumulates reduced ovothiol. **A.** (A) Extracted ion chromatograms of coumarin derivatives of ovothiol A, B, and C in Chlamydomonas cell extracts separated on a C18 reversed-phase column. (B) High-resolution mass spectrum of the ovothiol A-coumarin derivative. (C) Extracted ion chromatograms of ovothiol A-NEM (*N-*ethylmaleimide) derivative and ovothiol A disulfide from Chlamydomonas extracts separated on an amino normal-phase column. (D) High-resolution mass spectrum of the ovothiol A-NEM derivative. Unlabeled peaks originate from other compounds. In (A) and (C), all chromatograms start at time 0, but they are offset to the right at regular intervals to visually separate any peaks with identical retention times. In (B) and (D), only the *m/z* values corresponding to ovothiol A derivatives isotopes are represented. *m*/*z* calculated: ovothiol A-coumarin [M+H]^+^, 390.1118; ovothiol B-coumarin [M+H]^+^, 404.1275; ovothiol C-coumarin [M+H]^+^, 418.1431; ovothiol A-NEM [M+H]^+^, 327.1122; ovothiol A-disulfide [M+H]^+^, 401.1060.

In most cases, thiols react with ROS in their reduced form (Holler and Hopkins 1990; Marjanovic et al. 1995; Servillo et al. 2015; Osik et al. 2021; Hartmann et al. 2023). To determine the redox status of ovothiol A, the thiol group was blocked with *N*-ethylmaleimide (NEM; Supplementary Figure S1B) which prevents oxidation and disulfide bond formation following cell lysis. Under constant low light conditions (50 µmol photons m^-2^ s^-1^), only ovothiol A-NEM was detected by LC-MS, whereas ovothiol A disulfide was absent (Figures 3C and D). In Chlamydomonas, ovothiol A is therefore present in its reduced form, likely available to react with ROS.

### *OVOA1* mediates ovothiol A biosynthesis in Chlamydomonas

In bacteria, the biosynthesis of ovothiol A is catalyzed by two enzymes, OvoA and OvoB (Figure 1B). OvoA is a bifunctional enzyme responsible for the first and last steps: sulfoxide synthesis (Braunshausen and Seebeck 2011) and methylation of the imidazole ring (Naowarojna et al. 2018). OvoA enzymes are members of the non-heme iron sulfoxide/selenoxide synthase (NHISS) family, which are similar in sequence but can be differentiated based on specific functional residues (Kayrouz et al. 2024 and references within). The Chlamydomonas genome contains a single homologous sequence with the gene identifier Cre08.g380000_4532.1, which we tentatively named *OVOA1.* The N-terminal sulfoxide synthase (Figure 4A) and C-terminal methyltransferase (Figure 4B) domains of OVOA1 contain almost all the residues known for their catalytic activity or to bind cofactors, substrates, and products in OvoA homologs. OVOA1 also lacks functional residues specifically attributed to other NHISS, further supporting its possible role as an ovothiol synthase.

**Figure 4.**
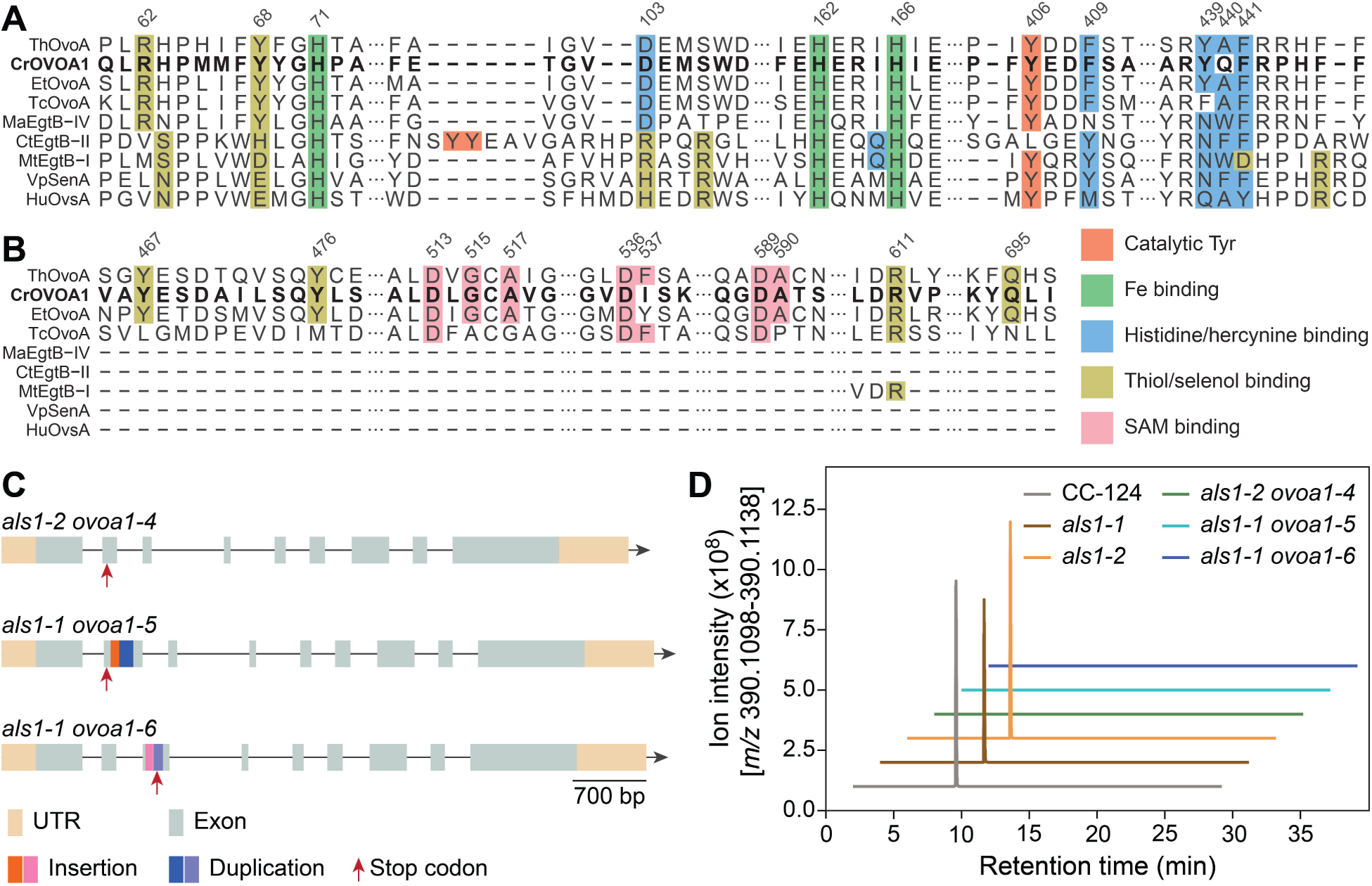
*OVOA1* mediates ovothiol A production in Chlamydomonas. Alignment of selected regions of the (A) sulfoxide synthase and (B) C-terminal methyltransferase domains of CrOVOA1 (Cre08.g380000_4532.1) with representative non-heme iron sulfoxide/selenoxide synthases. Previously characterized residues were colored by function according to Kayrouz et al. 2024, Ireland et al. 2025, and references within. See Supplementary Figure S3 for OVOA1 *in vitro* sulfoxide synthase activity measurement. EgtB, SenA, and OvsA are part of the biosynthetic pathways of ergothioneine (Stampfli et al. 2019), selenoneine (Liu et al. 2024), and ovoselenol (Kayrouz et al. 2024), respectively. Amino acids are numbered based on the ThOvoA sequence from *Hydrogenimonas thermophila* (Wang et al. 2023b). Et, *Erwinia tasmaniensis*; Tc, *Trypanosoma cruzi*; Ma, *Microcystis aeruginosa*; Ct, *Chloracidobacterium thermophilum*; Mt, *Mycolicibacterium thermoresistibile*; Vp, *Variovorax paradoxus*; Hu, *Halomonas utahensis*. (C) Schematic of *ovoa1* knockout mutants with premature stop codons, duplications of the flanking genomic DNA, and insertions of the homologous DNA template introduced in *OVOA1* by CRISPR-Cas9-mediated genome editing; these mutants are also edited in the selectable marker gene *ALS1* (Supplementary Dataset S1). UTR, untranslated region. (D) Extracted ion chromatograms of ovothiol A-coumarin from the *ovoa1* mutants and reference strains. Calculated *m/z* of ovothiol A-coumarin [M+H]^+^ = 390.1118.

To test the enzymatic function of OVOA1 from Chlamydomonas, the gene was tagged with a sequence for an N-terminal His tag and expressed in *E. coli*. The sulfoxide synthase activity of the partially purified protein was quantified *in vitro* using histidine and cysteine as substrates (Supplementary Figure S3). OVOA1 exhibited a high specific activity of 0.086 ± 0.004 µmol min^-1^ mg^-1^. Its observed turnover rate *k*_obs_ of 0.15 ± 0.01 s^-1^ is similar to that of OvoA from *Erwinia tasmaniensis* (Goncharenko et al. 2020), supporting the role of Chlamydomonas OVOA1 in ovothiol biosynthesis. The predicted methyltransferase activity of OVOA1 was not tested *in vitro*.

To examine the function of OVOA1 *in vivo*, we generated *ovoa1* knockout mutants using CRISPR-Cas9-mediated genome editing. We isolated three independent *ovoa1* mutants carrying early in-frame stop codons, insertions of the homologous DNA template, and/or duplications of flanking sequences which knock out the entire bifunctional gene (Figures 4C; Supplementary Figures S4 and S5; Supplementary Dataset S1). Because the method used is based on co-editing of *OVOA1* and the selectable marker *ALS1* (Akella et al. 2021), the *als1-1* and *als1-2* strains only edited in *ALS1* were generated and used as reference strains (Supplementary Table S2; Supplementary Dataset S1). These *als1* mutations had no effect on ovothiol A production (Figure 4D). Importantly, no ovothiol A-coumarin was detected in any of the *als1 ovoa1* CRISPR mutants (hereafter abbreviated as “*ovoa1*”; Figure 4D), which was additionally confirmed with two independent insertional mutants (*ovoa1-1* and *ovoa1-2*; Supplementary Figure S6). With these results, we conclude that *OVOA1* is required for ovothiol A biosynthesis in Chlamydomonas. In addition, the *OVOA1* transcript was expressed at a high basal level under low light growth conditions (within the top 15% of the most highly expressed genes; Supplementary Figure S7), which is consistent with the high concentration of ovothiol A that we measured. Finally, OVOA1 was predicted to be cytosolic by all softwares used: PredAlgo (Tardif et al. 2012), PB-Chlamy (Wang et al. 2023a), TPpred3 (Savojardo et al. 2015), TargetP2.0 (Armenteros et al. 2019), and DeepLoc2.1 (Ødum et al. 2024), suggesting that ovothiol A is synthesized in the cytosol.

### Ovothiol A biosynthesis is essential for singlet oxygen resistance

Since ovothiol A is a powerful antioxidant (Holler and Hopkins 1990; Marjanovic et al. 1995; Osik et al. 2021), we hypothesized that it could be involved in photooxidative stress responses in Chlamydomonas. In low light, the *ovoa1* mutants had no growth defect (Supplementary Figure S8), so we first tested if *OVOA1* was light-regulated by quantifying its transcript levels (Figure 5A). Upon high light exposure, the *OVOA1* mRNA level increased ∼ 3.5-fold within 15 min before stabilizing. Similar induction by light was observed in high-resolution expression data of synchronized cells during the diurnal cycle where *OVOA1* transcript levels spiked after the dark to light transition (Supplementary Figure S9; Zones et al. 2015; Strenkert et al. 2019). A moderate gradual increase was also visible before dawn, indicating that *OVOA1* may also be regulated by the circadian rhythm.

**Figure 5.**
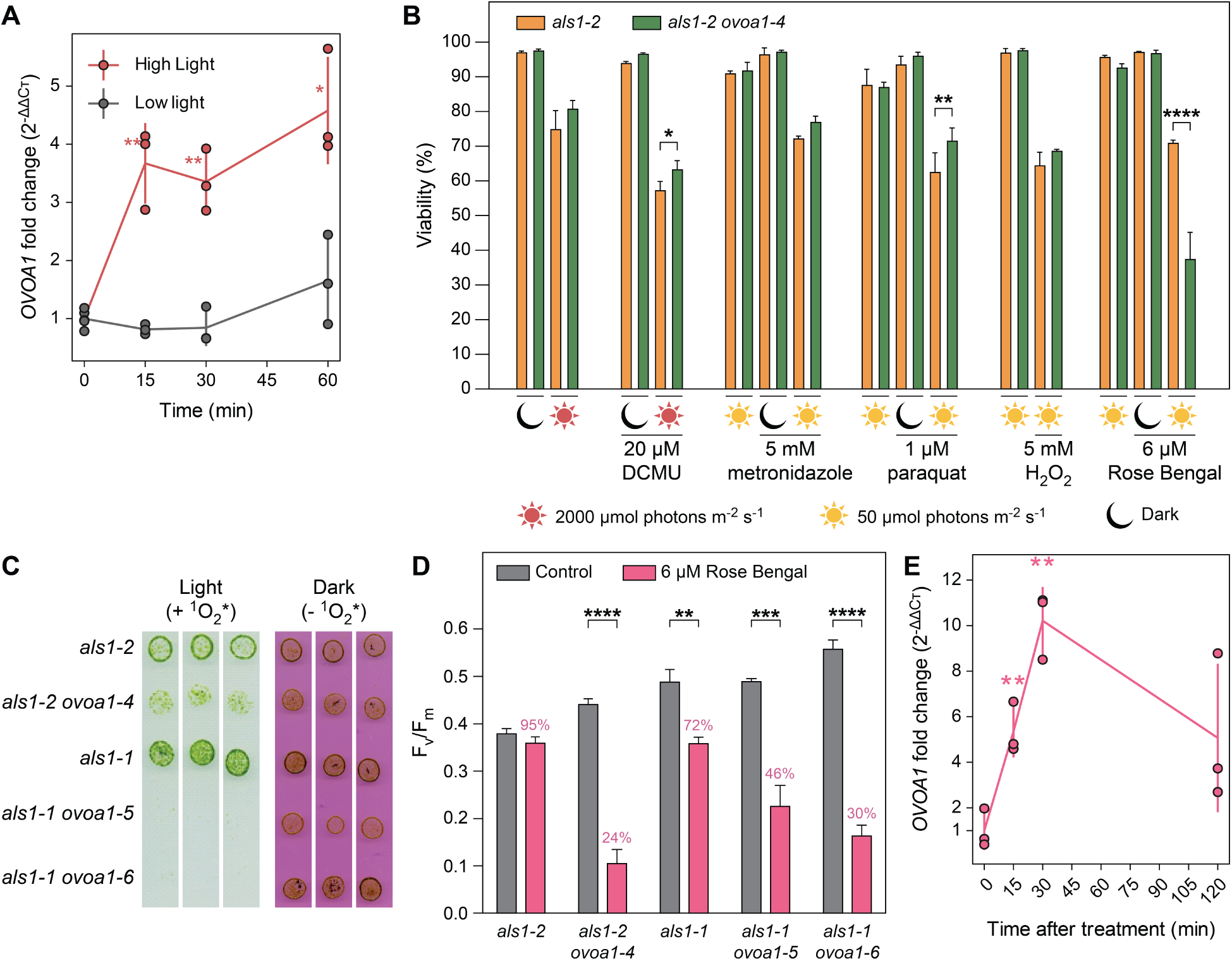
*OVOA1* is essential for singlet oxygen resistance. (A) *OVOA1* mRNA level under high (550 µmol photons m^-2^ s^-1^) or low (80 µmol photons m^-2^ s^-1^) light measured by RT-qPCR. (B) Sensitivity of the *ovoa1-4* mutant and reference strain to high light and reactive oxygen species. After treatments, cell viability was quantified with the CellTox Green Cytotoxicity Assay. (C) Growth of the *ovoa1* mutants and reference strains on TAP plates in the presence of ^1^O_2_* (after 4 days with 4 µM Rose Bengal in the light) or in the absence of ^1^O_2_* (after 8 days with 4 µM Rose Bengal in the dark). See Supplementary Figure S10 for the complete dataset. (D) Maximum quantum yield of photosystem II (F_v_/F_m_) of the *ovoa1* mutants and reference strains with or without exposure to 6 µM Rose Bengal for 6 h in the light. For each strain, the percentage corresponds to the fraction of F_v_/F_m_ in the Rose Bengal treatment compared to the control. (E) RT-qPCR measurement of *OVOA1* induction after treatment with 6 µM Rose Bengal in the light (80 µmol photons m^-2^ s^-1^). For (A) and (E), *t*-tests were used to compare each timepoint against the initial condition. For (B) and (D), one-way ANOVA with a Tukey-Kramer multiple comparison *post-hoc* test was used. In (B), only the comparisons between *ovoa1-4* and the reference strain are shown, other *p*-values are available in Supplementary Table S3. * *p* ≤ 0.05; ** *p* ≤ 0.01; *** *p* ≤ 0.001; **** *p* ≤ 0.0001. All experiments were done with at least three biological replicates and means ±SD are represented.

Based on these results and knowing that excess light induces ROS production in photosynthetic organisms, the *ovoa1-4* mutant was subjected to high light stress and other ROS-inducing treatments (Niemeyer et al. 2021). The exposure to high light (with or without DCMU), metronidazole, paraquat, or hydrogen peroxide significantly decreased the viability of the reference strain, but no additional defect was observed in *ovoa1-4* (Figure 5B). The latter showed reduced viability only when treated with Rose Bengal (Figure 5B), a photosensitizer that generates ^1^O_2_* in the light (Fischer et al. 2004). To confirm that the observed phenotype was due to the disruption of *OVOA1*, ^1^O_2_* sensitivity was tested with the two other CRISPR alleles and two insertional mutants. All mutants were significantly more sensitive to Rose Bengal in the light, indicating that *OVOA1* and ovothiol A are essential for ^1^O_2_* resistance in Chlamydomonas (Figure 5C; Supplementary Figure S11). The protective role of ovothiol A against ^1^O_2_* was further demonstrated by the significantly lower maximum efficiency of photosystem II (F_v_/F_m_) in the mutants compared to the reference strains after Rose Bengal treatment (Figure 5D). Finally, we found that *OVOA1* expression was induced after 15 min of exposure to 6 µM Rose Bengal under light, which reached a maximum of 10-fold induction after 30 min (Figure 5E). This result shows that, similarly to high light (Figure 5A) and the diurnal cycle (Supplementary Figure S9), ^1^O_2_* quickly induces *OVOA1* at the transcript level. We conclude that ovothiol A is essential for ^1^O_2_* resistance and is likely part of the early response to this ROS.

### Ovothiol A is involved in singlet oxygen acclimation and regulated by SAK1

In addition to being equipped with compounds that directly scavenge ^1^O_2_*, Chlamydomonas is able to acclimate to it: upon exposure to a sublethal dose of ^1^O_2_*, cells activate a specific set of genes that enable them to survive a subsequent higher dose (“challenge”) that would be lethal without pretreatment (Figure 6A; Ledford et al. 2007; Wakao et al. 2014). First, we found that ovothiol A concentration was not statistically changed after pretreatment, but it was reduced by more than half after the challenge, suggesting that it reacts with ^1^O_2_* (Figure 6B). The ability to acclimate to ^1^O_2_* was attenuated in all three *ovoa1* mutants compared to the reference strains. While pretreatment increased resistance to a moderate challenge in all strains, the mutants did not survive the higher concentrations of Rose Bengal, as was seen for the reference strains (Figure 6C). Furthermore, *OVOA1* gene expression was quickly induced, with a similar increase (∼10-fold) during pretreatment with 1 µM Rose Bengal as with higher doses of 6-8 µM (Figures 5E and 6D). These data suggest that *OVOA1* gene induction in response to a low non-lethal dose of ^1^O_2_* is one of the mechanisms that allow Chlamydomonas cells to resist more robustly a subsequent challenge.

**Figure 6.**
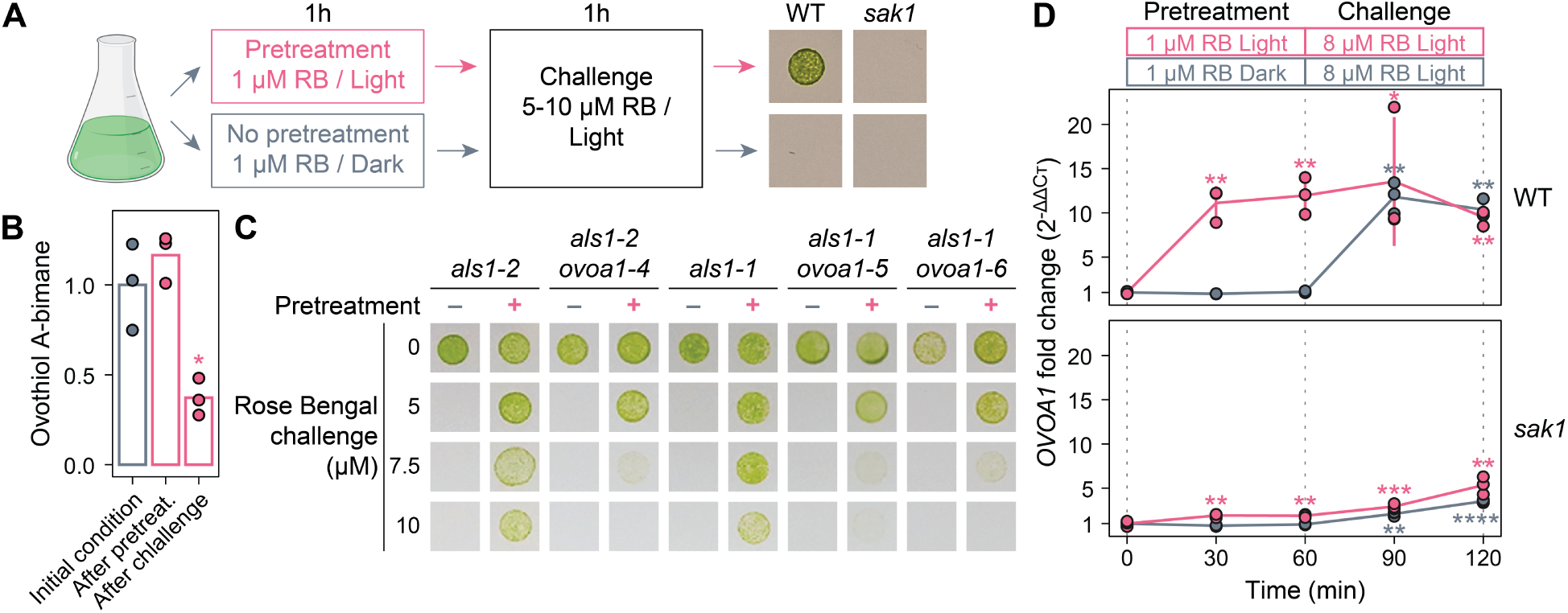
*OVOA1* is involved in singlet oxygen acclimation and regulated by SAK1. (A) Experimental design of ^1^O_2_* acclimation. Cells were pretreated with a sublethal dose of ^1^O_2_* generated by 1 µM Rose Bengal in the light. This “pretreatment” protects cells from a subsequent high dose (“challenge”, 5-10 µM Rose Bengal) that would be lethal without pretreatment. In the “no pretreatment” controls, cells were treated under the same condition but with pretreatment in the dark, which does not generate ^1^O_2_* (Ledford et al. 2007). (B) Ovothiol A-bimane derivative quantification in initial conditions, after 1 µM pretreatment, and after 8 µM challenge measured by LC-MS. (C) ^1^O_2_* acclimation of the *ovoa1* mutants and the reference strains. The cells were subjected to 5 to 10 µM challenge with or without pretreatment, and survival was monitored on plates after 4 days. (D) *OVOA1* expression measured by RT-qPCR in the 4A wild type and *sak1* (Wakao et al. 2014) during ^1^O_2_* acclimation with or without pretreatment. For (B) and (D), *t*-tests were used to compare each timepoint with the initial condition and *p*-value symbols are colored accordingly. * *p* ≤ 0.05; ** *p* ≤ 0.01; *** *p* ≤ 0.001; **** *p* ≤ 0.0001. All the experiments were done with three biological replicates and means ± SD are shown. RB, Rose Bengal.

The expression of *OVOA1* was then measured in *sak1*, a mutant disrupted in a phosphoprotein essential for ^1^O_2_*-induced cellular response and signaling (Wakao et al. 2014). In this mutant that does not survive the challenge after pretreatment (Figure 6A), *OVOA1* induction is almost completely abolished during pretreatment and reduced by half compared to the wild type after challenge (Figure 6D). This indicates that SAK1 is a major regulator of *OVOA1* expression during ^1^O_2_* acclimation and stress in Chlamydomonas. Nonetheless, in *sak1*, a weaker but statistically significant induction was observed during pretreatment and challenge, indicating that a SAK1-independent ^1^O_2_* signaling pathway exists, although *OVOA1* induction relies on it to a much lesser extent. Altogether, the ^1^O_2_*-induced decrease in ovothiol A, phenotype of the *ovoa1* mutants, as well as the expression of *OVOA1* in the wild type and in *sak1* demonstrate that ovothiol A biosynthesis is an essential protection mechanism activated during ^1^O_2_* acclimation.

### *OVOA1* is co-expressed with photooxidative stress response genes and is regulated by light and retrograde signaling pathways

To complement our characterization of the regulation of *OVOA1* and decipher its function in detail, we searched for co-expressed genes and pathways. Pearson correlation coefficients (PCCs) were calculated using 461 normalized RNA-seq datasets (Supplementary Figure S12A-C) of wild types collected in conditions related to photosynthesis and photooxidative stress (*e.g.* diurnal cycle, high light, carbon source, CO_2_ availability, and ROS) as well as other stress conditions (*e.g.* temperature, low copper, and anoxia). The top 150 genes co-expressed with *OVOA1* (*i.e.* with the highest PCCs) were then categorized (Figure 7A-C; Supplementary Figure S13C-D). This list was enriched in genes coding for predicted chloroplast proteins, which accounted for nearly 60% of the genes co-expressed with *OVOA1* (Figure 7A) and even more for larger PCCs (Supplementary Figure S13A). Strikingly, many of these genes were related to photosynthesis, photoprotection, and ROS responses (Figures 7B-C; Supplementary Figure S13B). *OVOA1* was highly co-expressed with the biosynthetic pathways of the carotenoids lutein and zeaxanthin as well as tocopherols and plastoquinol (Figure 7C), all of which are prominent ^1^O_2_* scavengers in plants and green algae (Li et al. 2012, 2016; Nowicka and Kruk 2012; Wakao and Niyogi 2021). In particular, the putative tocopherol methyltransferase gene *VTE6* (Wakao et al. 2021) had a high correlation coefficient (PCC = 0.73; Figure 7C; Supplementary Figure S12C). Other notable examples were genes involved in photosystem II repair, carbon fixation, or the CO_2_-concentrating mechanism (Figure 7C), such as *LCI16* and *EPYC1*, coding for components of the pyrenoid (Mackinder et al. 2016; Franklin et al. 2025). Additionally, genes coding for mitochondrial proteins were depleted among *OVOA1*-co-expressed genes compared to the whole genome (Figure 7A), and no gene related to mitochondrial oxidative stress was found (Figure 7B; Supplementary Figure S13B). These analyses showed that *OVOA1* is part of the transcriptomic response to photooxidative stress along with production of other antioxidants.

**Figure 7.**
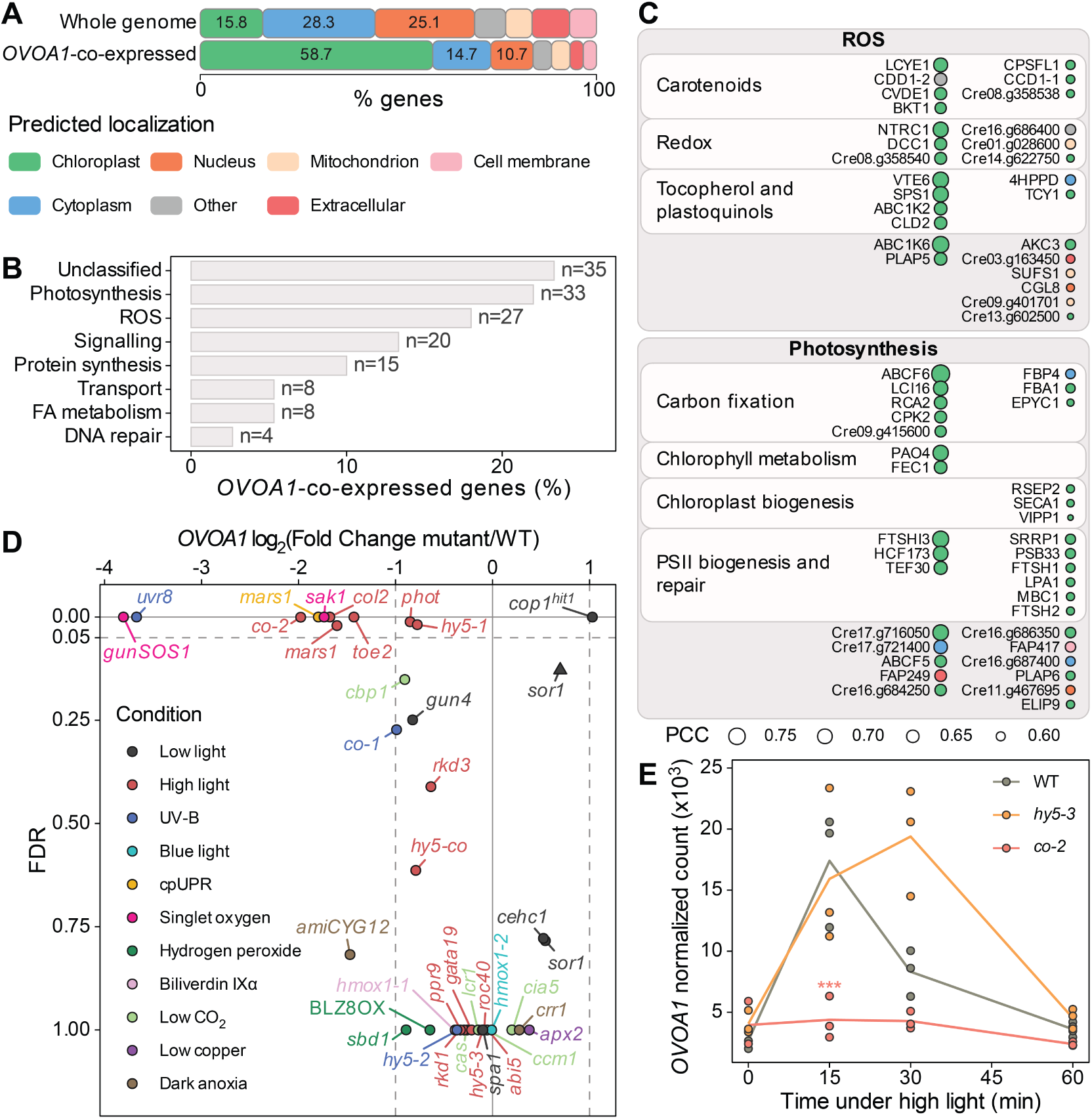
*OVOA1* is co-expressed with photoprotection genes and is regulated by singlet oxygen retrograde and light signaling pathways. (A-C) Analysis of the genes co-expressed with *OVOA1*. In normalized RNA-seq data, the 150 genes with the highest Pearson correlation coefficient (PCC) with *OVOA1* were selected (Supplementary Figures S12 and S13). (A) Predicted localization and (B) function of these co-expressed genes. For each functional category, “n” represents the number of genes which are described in detail in Supplementary Figure S13B and Supplementary Table S4. (C) Co-expressed genes in the “ROS” and “Photosynthesis” categories classified by predicted function. The localization color legend is shared between (A) and (C). (D) *OVOA1* expression in regulatory mutants. Using RNA-seq (circles) and microarray (triangle) transcriptomics, each mutant was compared with its reference wild type. Changes were considered significant if |log_2_(fold change)| ≥ 1 and false discovery rate (FDR) < 0.05 (dashed lines) (see Supplementary Table S5 and Material and methods for details and statistics). *gunSOS1*, *sak1*, and *mars1* are altered in ^1^O_2_* retrograde signaling; and *uvr8*, *phot*, *cop1^hit1^*, *toe, co*, *col2*, and *hy5* in light signaling. (E) *OVOA1* expression quantified by RNA-seq in *hy5-3*, *co-2*, and the wild type during high light exposure (350 μmol photons m⁻² s⁻¹). The points represent DESeq2-normalized counts, with three biological replicates per strain and timepoint. At each timepoint, Benjamini–Hochberg adjusted *p*-values (FDRs) were calculated with DESeq2 comparing each mutant with the wild type (*** *p* ≤ 0.001). BLZ8OX is a constitutive over-expressor of *BLZ8* and amiCYG12 is a strain where *CYG12* is downregulated by an artificial microRNA. FA, fatty acid; PSII, photosystem II; *gunSOS1*, *genomes uncoupled singlet oxygen signalling 1*; *sak1*, *singlet oxygen acclimation knocked-out 1*; *mars1*, *mutant affected in chloroplast-to-nucleus retrograde signaling 1*; *uvr8*, *uv resistance locus 8*; *phot*, *phototropin*; *cop1^hit1^*, *constitutive photomorphogenic 1* / *high light tolerant 1*; *toe2*, *target of eat 2*; *co*, *constans*; *col2*, *constans-like 2*; *hy5*, *long hypocotyl 5*.

Because we found that *OVOA1* was induced by ^1^O_2_* and light (Figures 5 and 6; Supplementary Figure S9), we sought to characterize its regulation in more detail. We re-analyzed transcriptomic data generated with mutants of 31 genes coding for components of cell signaling. These regulators are involved in photoprotection and other processes like the CO_2_-concentrating mechanism, anoxia, or copper starvation. Each mutant was compared with the relevant reference strain in the reported condition to calculate the *OVOA1* expression fold change using DESeq2 methods (Love et al. 2014) for normalization, differential expression, and statistics (Figure 7D; Supplementary Table S5). The most striking effect on *OVOA1* was observed in mutants impaired in retrograde signaling induced by ^1^O_2_* (*gunSOS1* and *sak1*) and the chloroplast unfolded protein response (*mars1*) as well as in signaling components of light stress responses (*uvr8, cop1^hit1^*, *toe2*, and *col2*) (Youssef et al. 2023; Wakao et al. 2014; Perlaza et al. 2019; Tilbrook et al. 2016; Gabilly et al. 2019; Tokutsu et al. 2019; Wulf et al. 2023). For *sak1*, a similar downregulation of *OVOA1* transcript level was found in this RNA-seq analysis (Figure 7D) and by RT-qPCR (Figure 6D), strengthening our results. The *phot* mutant, which is defective in the blue light photoreceptor phototropin, had a moderate but statistically significant effect on *OVOA1* expression, indicating a possible involvement of the PHOT-DDB1/DET1 pathway (Aihara et al. 2019). A direct regulation by light is also supported by the high co-expression of *OVOA1* with the genes coding for RUP1 (a regulator of the UV-B photoreceptor UVR8) and the photoreceptor DCRY1 (Supplementary Figure S13B; Tilbrook et al. 2016; Rredhi et al. 2024).

To complement these analyses, we then performed a high light RNA-seq time course for *co-2* (=*constans*) and *hy5-3*, mutants defective in two putative COP1-dependent transcription factors known to regulate photoprotection in Chlamydomonas (Gabilly et al. 2019; Lämmermann et al. 2020; Wulf et al. 2023; Benko et al.). The strong and rapid induction (within 15 min) of *OVOA1* was retained in *hy5-3*, whereas it was completely abolished in *co-2* (Figure 7E). Finally, it caught our attention that *OVOA1* expression is dependent on the ^1^O_2_* response regulators SAK1 and gunSOS1/TSPP1 (Figures 6D and 7D), while mutants of regulators involved in the response to hydrogen peroxide, like BLZ8 and SBD1, did not show a significant effect (Choi et al. 2022; Koletti et al. 2022). This is in agreement with our finding that *OVOA1* has a role specific to ^1^O_2_* and not to other ROS (Figure 5B).

Taken together, our results show that ovothiol A biosynthesis is regulated by both retrograde and light-induced signaling pathways in which SAK1, gunSOS1, MARS1, UVR8, and CO were identified as major regulators. The working model presented in Figure 8 summarizes *OVOA1* regulation based on our results and known signaling pathways.

**Figure 8.**
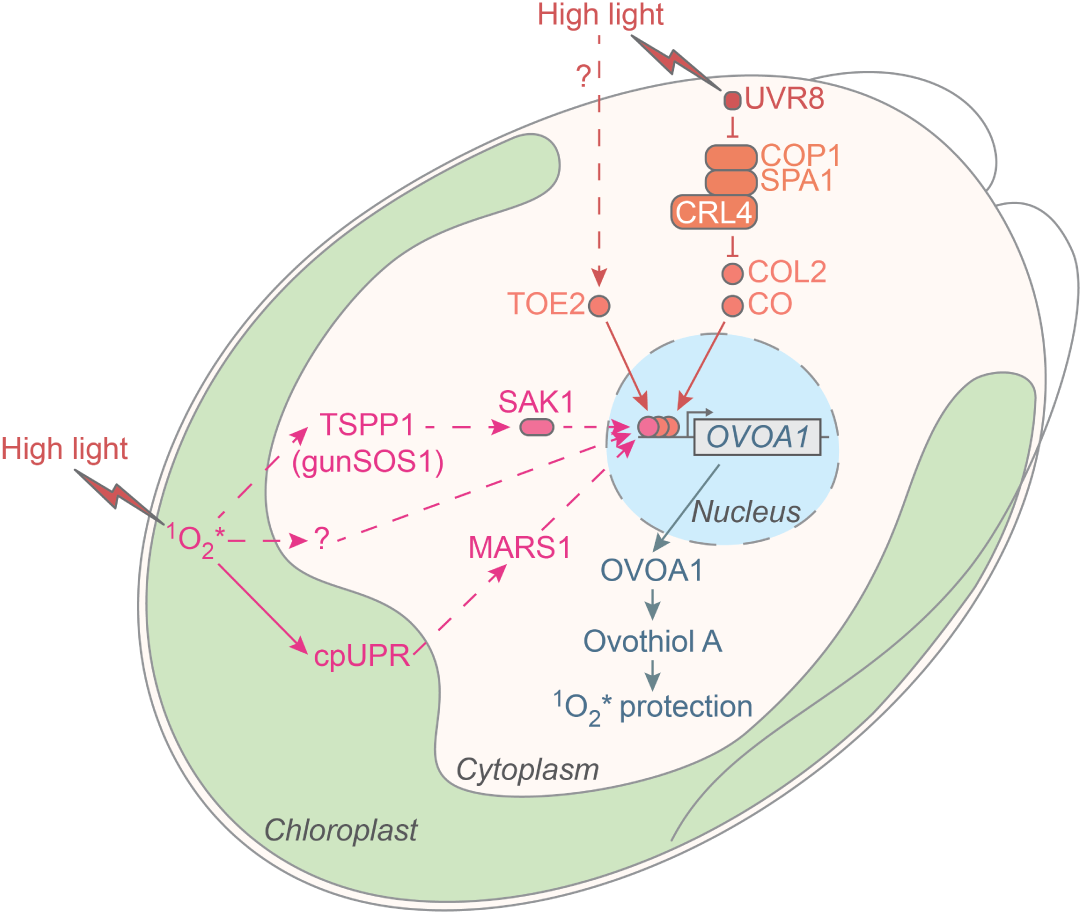
Model of ovothiol A function and regulation in Chlamydomonas. High light generates ^1^O_2_* in the chloroplast, and *OVOA1* is activated by retrograde signals (in pink) mediated by gunSOS1/TSPP1 and SAK1 (Youssef et al. 2023; Wakao et al. 2014) and the MARS1 kinase-dependent chloroplast unfolded protein response (cpUPR) (Perlaza et al. 2019). A *sak1*-independent moderate activation of *OVOA1* (Figure 6D) could be mediated by an unknown pathway (pink question mark) or the MARS1 pathway. *OVOA1* is also activated through light signaling pathways (in orange) dependent on the photoreceptor URV8, the COP1/SPA1 E3 ubiquitin ligase complex, and the transcription factors COL2 and CO (Tilbrook et al. 2016; Gabilly et al. 2019; Tokutsu et al. 2019; Lämmermann et al. 2020; Wulf et al. 2023). COL2, CO, and TOE2 are master regulators of photosynthesis and photoprotection in Chlamydomonas (*e.g.* non-photochemical quenching). TOE2 is regulated by blue light in Arabidopsis but remains poorly characterized in Chlamydomonas (Wulf et al. 2023). Finally, these transcription factors activate *OVOA1*, which mediates ^1^O_2_* resistance and acclimation (in blue). Solid and dashed lines represent effects speculated to be direct or indirect, respectively. TSPP1, trehalose 6-phosphate phosphatase 1; CRL4, cullin-RING ligase 4 subunit; see legend for Figure 7 for other abbreviations.

## Discussion

The response of plants and unicellular algae to oxidative stress has been extensively investigated, including numerous studies on the role of antioxidants such as glutathione (reviewed in Zaffagnini et al. 2012; Wakao and Niyogi 2021). Strikingly, the role of other thiol antioxidants like ovothiols, has been completely overlooked. In this study, we demonstrate that ovothiol A is a major antioxidant in the model green alga Chlamydomonas (Figure 3). Furthermore, we show that *OVOA1* is essential for ovothiol A biosynthesis as well as for ^1^O_2_* resistance and acclimation (Figures 4, 5, and 6). Finally, *OVOA1* is found to be regulated by both ^1^O_2_* retrograde signaling and light response pathways (Figures 6, 7 and 8).

In diatoms, the low ovothiol B concentration of 50 µM suggests a minor antioxidant or regulatory role (Milito et al. 2020a, 2020b; Russo et al. 2023). It has been proposed that ovothiol A disulfide regulates plastid ATP synthase activity in the green alga *Dunaliella salina* (Selman-Reimer et al. 1991), and similar regulatory functions are also possible in Chlamydomonas. However, we favor the primary function of ovothiol A in Chlamydomonas as an antioxidant given the millimolar concentration in the cells. Such cellular concentration is comparable or higher than ovothiol production in other eukaryotes like sea urchin eggs or trypanosomatids (Turner et al. 1988; Ariyanayagam and Fairlamb 2001). This is also a typical concentration of a related antioxidant, ergothioneine, detected in bacteria and eukaryotes (Hartman 1990; Dumitrescu et al. 2022). Moreover, a high concentration indicates that ovothiol A is probably a final product and not an intermediate used to produce another metabolite (Castellano and Seebeck 2018). Strikingly, at 2.3 mM, ovothiol A is even more abundant than glutathione, which is reported around 1 mM in Chlamydomonas (Stoiber et al. 2007).

Being crucial for plants, oxidative stress defenses are often redundant. For example, tocopherol over-production can compensate for the lack of zeaxanthin and lutein in Chlamydomonas (Li et al. 2012). This raises the question whether ovothiol A has a different function than other ^1^O_2_* scavengers. *OVOA1* is highly co-expressed with tocopherols, plastoquinol, lutein, and zeaxanthin biosynthesis genes (Figure 7C). On the other hand, these isoprenoids are lipid-soluble and produced in the chloroplast (Krieger-Liszkay et al. 2008), whereas ovothiol A is water-soluble and likely cytosolic based on the predicted localization of OVOA1. These results suggest that all these antioxidants are acting in synergy in different subcellular compartments. The existence of parallel mechanisms of photoprotection, antioxidants, and non-photochemical quenching (NPQ), could explain why the *ovoa1* mutants do not show high light sensitivity in our experiments (Figure 5B). It will be interesting to test whether the viability of the mutants is impaired when high light is combined with other types of stress such as elevated temperatures or low CO_2_ availability.

The regulation of *OVOA1* by retrograde signaling, light, and associated signaling pathways (Figure 8) suggests that ovothiol A is involved in the photooxidative stress response. *OVOA1* activation by high light is weaker than that of the NPQ genes *LHCSR1*, *PSBS1*, and *PSBS2*, which are induced more than 150-fold (Gabilly et al. 2019). This is probably because, under low light conditions, *OVOA1* is expressed at a much higher basal level than these NPQ genes (Supplementary Figure S7). Nevertheless, and most importantly, *OVOA1* and other photoprotection genes like *LHCSR1*, *PSBS1*, *PSBS2*, and *LHL4* (Dannay et al. 2025) share common activation pathways mediated by UVR8-COP1-CO, COL2, TOE2, and possibly PHOT-DDB1/DET1. This shows that the targets of these light response pathways are more diverse than previously thought in Chlamydomonas. Regulation of some nuclear genes by both photoreceptor-driven pathways and chloroplast-to-nucleus retrograde signaling has been observed in plants and Chlamydomonas (Richter et al. 2023). For example, *LHCSR1* is activated by the UVR8-COP1-CO pathway (Gabilly et al. 2019) as well as SAK1- and GUN4-mediated retrograde signaling (Wakao et al. 2014; Formighieri et al. 2012) but is independent of MARS1 (Perlaza et al. 2019). The activation of *OVOA1* by multiple pathways (Figure 8) is therefore another remarkable example of converging regulation, highlighting that environmental cues dynamically regulate *OVOA1* despite a high basal level of expression.

In nature, intracellular ^1^O_2_* is not only generated in the presence of high light, but also in response to other stresses such as increased temperature (Krieger-Liszkay et al. 2008; Prasad et al. 2016). In addition, ^1^O_2_* can originate from the environment (*e.g.* in soil; reviewed in Larson and Marley 1999). We and others have described the regulation of ^1^O_2_* acclimation by SAK1, gunSOS1/TSPP1, and SOR1 (Wakao et al. 2014; Youssef et al. 2023; Fischer et al. 2012). However, besides the possible involvement of GPX5/GPXH and GSTS1, the defense mechanisms mediating cell survival itself during ^1^O_2_* acclimation remained to be identified. For example, antioxidant carotenoids and α-tocopherol are not involved in the acclimation process (Ledford et al. 2007). Here, we show that ovothiol A is one of the induced defense mechanisms that makes cells more resistant to ^1^O_2_* during acclimation. However, *ovoa1* mutants are only partially deficient in acclimation (Figure 6C), indicating that other mechanisms are also involved in this process.

To the best of our knowledge, our study is the first description of ovothiol being involved in resistance against ^1^O_2_*. The reaction of ^1^O_2_* with ovothiols remains to be demonstrated experimentally, but it has been well characterized for ergothioneine, another histidine-derived thiol (Hartmann et al. 2023). Given the structural similarity of ovothiols and ergothioneine, we propose the following model: ^1^O_2_* may react with one of the carbons in the ovothiol imidazole ring to form a hydroperoxide that may then be reduced by glutathione. On its own, glutathione is a poor scavenger of ^1^O_2_* (Devasagayam et al. 1991), so we hypothesize that ovothiol A is essential by reacting with ^1^O_2_* directly. Ovothiol reacts rapidly with hydrogen peroxide (Turner et al. 1988; Osik et al. 2021) but, in Chlamydomonas, multiple scavengers are known for this ROS such as glutathione and ascorbate in combination with glutathione and ascorbate peroxidases (Wakao and Niyogi 2021). This makes ovothiol A less likely to be essential for hydrogen peroxide resistance, which is supported by the fact that the *ovoa1* mutants are more sensitive to ^1^O_2_* but not to hydrogen peroxide compared to the reference strain (Figure 5B). A similar mechanism was hypothesized in trypanosomatids: the presence of trypanothione and glutathione peroxidases diminishes the importance of ovothiol to detoxify hydrogen peroxide (Ariyanayagam and Fairlamb 2001). Lastly, crosstalk between ROS and reactive electrophile species has been described (Fischer et al. 2012; Wakao and Niyogi 2021), and the OVOA1 protein is induced more than 50-fold in response to a nitrosative stress (Lambert et al. 2024). This suggests that ovothiol A has broad functions, and studying its role in response to all these stresses offers exciting avenues for future research.

The fact that the importance of ovothiol A went unnoticed for decades in a model organism such as Chlamydomonas highlights that major aspects of the function, diversity, and evolution of small thiols remain to be discovered. Furthermore, we show that a wide diversity of ecologically relevant microalgae possesses OvoA homologs and are likely producing ovothiols (Figure 2). Prominent examples include diatoms, coccolithophores, and the coral symbionts *Symbiodinium*, which are major contributors to carbon fixation and sequestration of CO_2_ into biominerals (Guidi et al. 2016; Ziveri et al. 2025). Small thiols may therefore play a more important role than previously thought in microalgal ecology and global biochemical cycles.

## Materials and methods

### Strains and culture conditions

The wild-type strain CC-124 of *C. reinhardtii* was used, except for experiments with *sak1* (CC-5795), *spa1*, *hy5-3*, or *co-2*, for which the reference strain was 4A (CC-4051; Wakao et al. 2014; Gabilly et al. 2019), and for experiments with the CLiP mutant LMJ.RY0402.136602, for which the reference strain was CC-4533 (Li et al. 2019). These strains were previously generated by us or obtained from the Chlamydomonas Resource Center (www.chlamycollection.org) and are summarized in Supplementary Table S2. Unless otherwise specified, cells were grown in Tris-acetate-phosphate (TAP) medium (Hui et al. 2023) at 20 to 25 °C under continuous white light (50 to 100 µmol photons m^-2^ s^-1^) and shaking.

### Identification of ovothiol A

CC-124, the *als1* reference strains, and the *ovoa1* mutants were cultivated for 5 days to reach early stationary phase. An aliquot containing 3 x 10^8^ cells was centrifuged at 4,500 x *g* at 4 °C for 5 min and resuspended in 300 µL of sodium acetate buffer (40 mM sodium acetate pH 4.6, 2 mM EDTA pH 8.0, 1 mM tris-2-carboxyethyl-phosphine (TCEP)). The suspension was then extracted with 300 µL of chloroform. 270 µL of the aqueous phase were collected and derivatized with 30 µL of 10 mM 4-bromomethyl-7-methoxycoumarin (BMC, prepared in DMSO) at 70 °C for 30 min in the dark. The reaction was stopped by adding 30 µL of formic acid to the derivatized sample, followed by centrifugation at 16,000 x *g*, 20 °C for 2 min. Ovothiol A-coumarin was analyzed with a Vanquish Horizon UHPLC coupled to an Orbitrap Q Exactive Plus mass spectrometer (Thermo Fisher Scientific) via a heated electrospray ion source (HESI). 1 µL of sample was injected and separated on a Kinetex C18 column (100 x 2.1 mm; 1.7 µm, Phenomenex) at 25 °C using a flow rate of 0.25 mL/min. Eluent A consisted of H_2_O/0.05% (v/v) formic acid, and eluent B was acetonitrile. After 5% B for 5 min, the composition of the mobile phase changed linearly to 100% B over the course of 10 min, and was kept at 100% B for 7 min. Over the last 3 min, the mobile phase changed back to 5% B for column equilibration before the next injection. The HESI source was operated in positive mode at 3500 V, and the temperature of the ion transfer tube was 300 °C. The resolution of the Orbitrap was set to 70,000 with a mass range of *m/z* from 150 to 2000 for BMC derivatives. The extracted ion chromatograms and mass spectra were processed with FreeStyle 1.8 SP2 QF1 (Thermo Fisher Scientific).

### Absolute quantification of ovothiol A

CC-124 cell extracts and L-2-thiohistidine standard solutions were prepared in 300 μL of sodium acetate buffer, extracted in chloroform, derivatized with BMC, and the reactions quenched with formic acid as described above. After filtration with a pore size of 0.45 μm, 10 μL of filtrate were analyzed using a JASCO HPLC 900 (JASCO Germany GmbH) equipped with a MD-910 diode array detector. A C18 column (Kromasil C18 HPLC column, 5 μm 4.6 x 150 mm, K08670356, Sigma-Aldrich) was equilibrated and used at 20 °C with a flow rate of 0.6 mL/min. Eluent A was 0.1% formic acid in water and eluent B was 0.1% formic acid in acetonitrile. The composition of the mobile phase changed linearly from 2% B to 100% B over the course of 15 min, was kept at 100% B for 5 min, changed back to 2% B over 1 min and kept at 2% B for 6 min before the next injection. The absorbance of BMC derivatives was quantified at 330 nm. The ovothiol A-coumarin peak was identified by comparing the chromatogram with the chromatogram obtained from a mutant lacking ovothiol A. The intracellular concentration of ovothiol A was calculated by assuming an intracellular volume of 51.6 fL (Schötz et al. 1972).

### Relative quantification of ovothiol A

For relative quantification of small thiols, mBBr (monobromobimane) derivatization was used (Newton and Fahey 1995; O’Neill et al. 2015). 1.5 x 10^7^ cells were resuspended in 477.5 µL of HEPES-TCEP-ACN buffer (2 mM HEPES, 2 mM TCEP in 50% acetonitrile). The mixture was incubated at 60 °C, 1400 rpm for 8 min, frozen in liquid nitrogen, incubated at 60 °C, 1400 rpm for 4 min, and lysed by bead beating with Lysing Matrix D in a FastPrep-24 5G (MP Biomedicals) with 3 cycles of 30 s at 6 m/s. Cell lysis was completed by freezing the mixture in liquid nitrogen, incubating it at 60 °C, 1100 rpm for 8 min, and repeating the bead beating. After spiking *N*-acetylcysteine amide (internal standard, 10 µM final concentration) in 420 µL of lysate, thiols were derivatized with 2 mM mBBr at 60 °C, 1400 rpm for 30 min in the dark. After cooling on ice, the reaction was stopped with 2.2 µL of 5 M methanesulfonic acid. The mixture was cleared of insoluble debris by centrifugation and filtering before being diluted 10-fold in water. LC-MS was performed by the QB3/Chemistry mass spectrometry facility at UC Berkeley. 10 µL were injected in an Agilent 1200 HPLC equipment (Agilent Technology) using an ZORBAX 300SB-C18 column (150 x 2.1 mm, 5 µm, Agilent). Phase A contained 0.1% formic acid in water and phase B 0.1% formic acid in 50% acetonitrile / 50% methanol. The following gradient was used at 40 °C with a flow rate of 0.3 mL/min: 5% B for 1 min, linear increase from 5% to 95% B in 6 min, and 95% B for 7 min. The phases were returned to 5% B in 1 min and the column was re-equilibrated for 15 min. High resolution mass spectrometry analysis was performed on an LTQ Orbitrap XL mass spectrometer equipped with an electrospray ionization source (Thermo Fisher Scientific) operated in positive ion and full scan mode with a mass range from *m/z* 100 to 700. The ovothiol A-bimane peak identity was confirmed with the *ovoa1-4* mutant compared with its reference strain *als1-2*.

### Measurement of ovothiol A redox state

CC-124 was cultivated for 5 days to reach early stationary phase, 3 x 10^8^ cells were centrifuged at 4,500 x *g* at 4 °C for 5 min and resuspended in 300 µl of 40 mM sodium acetate pH 4.6, 2 mM EDTA pH 8.0, 30 mM *N*-ethylmaleimide (NEM). The suspension was extracted with 300 µL of chloroform. NEM derivatives were quantified using the Q Exactive Plus LC-MS device described above, with a detection mass range of *m/z* from 65 to 975. 10 µL of sample were injected and separated on a Luna-NH_2_ column (150 x 2 mm, 3 µm, Phenomenex) at 25 °C using a constant flow rate of 0.15 mL/min. Eluent A consisted of 20 mM ammonium acetate pH 9.0, and eluent B was acetonitrile. The initial composition of 25% A linearly increased to 100% A over 10 min and was kept at 100% A for another 5 min. Over the next 5 min, the mobile phase changed back to 25% A and was kept at 25% A for another 5 min before the next injection.

### Heterologous expression and purification of OVOA1

*E. coli* BL21(DE3) harboring pL1SL2 encoding a chaperone of *Streptomyces* (Betancor et al. 2008) was transformed with a pET28 vector encoding His-tagged OVOA1 from Chlamydomonas (Cre08.g380000_4532.1.p; protein sequence provided in Supplementary Dataset S2). The TB-M-2LAC-SUC auto-induction medium used for the heterologous expression of OVOA1 contained 24 g L^−1^ yeast extract, 12 g L^−1^ tryptone, 1.0% glycerol, 0.1% glucose, 0.4% α-lactose monohydrate, 25 mM Na_2_HPO_4_, 50 mM NH_4_Cl, 5 mM Na_2_SO_4_, 25 mM KH_2_PO_4_, 10 mM Na_2_HPO_4_, 25 mM succinic acid, 10 µM FeCl_3_, 4 µM CaCl_2_, 2 µM MnCl_2_, 2 µM ZnSO_4_, 0.4 µM CoCl_2_, 0.4 µM CuCl_2_, 0.4 µM NiCl_2_, 0.4 µM Na_2_MoO_4_, 0.4 µM Na_2_SeO_3_, and 0.4 µM H_3_BO_3_ (Studier 2005). In a 3-L flask, 600 ml of TB-M-2LAC-SUC medium supplemented with 100 μg/mL of ampicillin and kanamycin was inoculated with 1 mL of overnight culture in LB medium. The culture was grown at 37 °C for 3 h with 220 rpm orbital shaking, after which the culture incubated for another 66 h at 20 °C. The cells were collected by centrifugation and frozen until purification. The frozen cell pellet was resuspended in washing buffer (50 mM Na phosphate, pH 8.0, 300 mM NaCl, and 30 mM imidazole) supplemented with 20 μg/mL lysozyme, 20 μg/mL DNase, 0.01% Triton X-100, and 1 mM MgCl_2_. The cells were lysed by sonication (Q500 sonicator, QSONICA) for 5 min at 50% amplitude with 2 s pulse intervals while being cooled. The lysate was centrifuged at 66,000 x *g* at 4 °C for 30 mins to remove the cell debris. The His-tagged OVOA1 protein was purified with Ni-NTA affinity beads (ROTI Garose, Carl Roth). After the applying the cleared lysate to the gravity column, the column was washed with washing buffer until the flowthrough showed a stable absorbance at 280 nm. Bound protein was eluted with elution buffer (50 mM Na phosphate, pH 8.0, 300 mM NaCl, and 250 mM imidazole). The purified protein was dialyzed against 50 mM Na phosphate, pH 8.0. After quantification of the absorbance at 280 nm using Nanodrop (Thermo Fisher), the protein concentration was calculated with the help of the extinction coefficient (Ɛ_280_ = 164,835 M^-1^ cm^-1^) predicted by ProtParam (Expasy).

### Quantification of sulfoxide synthase activity

The activity assay was adapted from a prior publication (Mashabela and Seebeck 2013). The reaction mixture comprised 1 mM L-histidine, 1 mM L-cysteine, 20 μM FeSO_4_, 1 mM ascorbate, 2 mM TCEP, 1 μM OVOA1, and 20 mM NaCl in 50 mM Na phosphate, pH 8.0. The 200-μL reaction mixture in a 1.5-mL tube was incubated at 25 °C with shaking at 600 rpm. Aliquots of 15 μL were removed at different times and quenched with 15 μL of 2% trichloroacetic acid. The consumption of histidine and the production of 5-histidylcysteine sulfoxide were monitored by ion-exchange HPLC with UV detection at 220 nm using a Shimadzu LC-40 series HPLC with a diode array detector and a Luna SCX 100 Å cation exchange column (150 x 4.6 mm, 5 μm, Phenomenex). The samples were eluted with a mixture of 20 mM phosphoric acid, pH 2.0 (eluent A) and 20 mM phosphoric acid, pH 2.0, 1 M NaCl (eluent B) at flow rate of 1 mL/min. After equilibration with 5% B, a linear gradient changed from 5% to 20% B over the course of 7 min followed by an increase to 99% B over the course of 0.5 min. The column was washed and equilibrated back to 5% B before measurement of the next sample.

The identity of the reaction product was verified by high-resolution mass spectrometry using a Shimadzu Nexera-X2 HPLC coupled with a Bruker maXis II ESI-TOF mass spectrometer. The HPLC was equipped with a Luna Omega Polar C18 100Å reversed-phase column (100 x 2.1 mm, 1.6 µm, Phenomenex). The samples were eluted with a mixture of water with 0.1% formic acid (eluent A) and acetonitrile with 0.1% formic acid (eluent B) at flow rate of 0.4 mL/min. After equilibration with 5% B for 1 min, a linear gradient changed from 5% to 95% B over the course of 2 min. The column was washed and equilibrated back to 5% B before measurement of the next sample. The mass spectra were internally calibrated using Na-formate solution prior to analysis.

### Generation of *ovoa1* mutants

The *ovoa1-4*, *ovoa1-5*, and *ovoa1-6* mutants were generated by CRISPR-Cas9-mediated genome editing based on the method of Akella et al. 2021. The design of crRNAs and single-stranded oligodeoxynucleotides (ssODNs), the assembly of the ribonucleoprotein complexes, their mixing with the ssODNs and the electroporation were performed as described (Lihanova et al. 2025; Supplementary Dataset S1; Supplementary Table S6). 33 transformants were screened by colony PCR as follows. Single colonies were resuspended in 100 µL of 10 mM EDTA pH 8.0 and lysed for 15 min at 100 °C in a thermocycler. After centrifugation at 16,000 x *g* for 2 min, the supernatant was diluted 3-fold and 2 µL were used in a 25-µL PCR reaction with the Q5 High-Fidelity DNA polymerase (NEB) targeting *ALS1* or *OVOA1*. For *OVOA1*, PCR products were further analyzed by restriction digestion (Supplementary Figure S4) and sequenced by Sanger sequencing. Primer sequences are provided in Supplementary Table S7.

To isolate the *ovoa1-1* mutant, a library of approximately 25,000 insertional mutants was generated with the wild type D66 (CC-4425) and screened by PCR as described in Pootakham et al. 2010 and Gonzalez-Ballester et al. 2011. The mutant CRMS104 edited in *OVOA1* was successively backcrossed with wild types CC-124, CC-125, and CC-124 again as described in Heimerl et al. 2018, yielding *ovoa1-1* (Supplementary Figure S6A). The *ovoa1-2* mutant was derived from the CLiP mutant LMJ.RY0402.159407 (abbreviated as “LMJ407” here; Li et al. 2019). After both borders of the insertion cassette (Supplementary Figure S6A) were confirmed by PCR and DNA sequencing, LMJ407 was successively backcrossed with CC-125 and CC-124, yielding the strain LMJ407-2.1 (=*ovoa1-2*).

### Viability measurements with high light and ROS treatments

The strains were grown for 3 days to reach the exponential growth phase, using TAP medium for the ROS treatments and TP medium (Hui et al. 2023) for the experiments with high light. The cell density was adjusted to 2 x 10^6^ cells/mL before the cells were treated with high light (2000 µmol photons m^-2^ s^-1^) with or without 20 µM 3-(3,4-dichlorophenyl)-1,1-dimethylurea (DCMU) for 48 h. The treatments with 6 µM Rose Bengal for 6 h, 5 mM metronidazole for 24 h, and 1 µM paraquat for 24 h were carried out both in low light (50 µmol photons m^-2^ s^-1^) and in the dark while the treatment with 5 mM H_2_O_2_ for 24 h was done in low light only. To quantify cell viability, 50 µL of cell suspension were pipetted into a well of a black 96-well plate and mixed with 50 µL of 2x CellTox Green Reagent, which was prepared by diluting the CellTox Green Dye 500 times with the assay buffer provided in the CellTox Green Cytotoxicity Assay kit (Promega). After incubation in the dark for 15 min and orbital shaking at 500 rpm for 1 min, the fluorescence signal was measured using a Spark microplate reader (Tecan), with excitation at 485 nm and a 530/20 nm band pass filter to detect emission. A reference sample with 100% cell death was prepared by killing cells at 100 °C for 30 min.

### Growth assay with Rose Bengal and singlet oxygen acclimation

The *als1-1* and *als1-*2 reference strains and the *ovoa1* mutants were grown for 5 days, the concentration was adjusted to 2 x 10^6^ cells/mL, and 10 µL were spotted onto TAP agar plates supplemented with Rose Bengal (2, 4, 6, or 8 µM) and grown in low light and in the dark for up to 8 days. Singlet oxygen acclimation was analyzed as previously described (Wakao et al. 2014). Briefly, *ovoa1* mutants, *als1-1*, *als1-2*, 4A, and *sak1* saturated cultures (15 x 10^6^ cells/ml) where transferred to 24-well plates with 1 mL per well and acclimated to the light conditions (80 µmol photons m^-2^ s^-1^) for 5 h. Cells were then pretreated with 1 µM Rose Bengal for 1 h in the light or in the dark before being challenged with 5.0, 7.5, 8.0, or 10.0 µM Rose Bengal for 1 h in the light. For 4A and *sak1*, at each timepoint, 1 ml was harvested by centrifugation and immediately flash-frozen in liquid nitrogen for RT-qPCR and LC-MS analyses. At the end of the treatment, drops of 3 µL were spotted onto TAP agar plates and cell survival was observed after a few days.

### RNA purification and RT-qPCR

4A cultures were prepared and light acclimated as for the ^1^O_2_* acclimation. The high light timecourse was performed at 550 µmol photons m^-2^ s^-1^ and the Rose Bengal timecourse with 6 µM Rose Bengal at 80 µmol photons m^-2^ s^-1^. At each timepoint, 1 mL was harvested by centrifugation and immediately flash-frozen in liquid nitrogen. RNA was extracted with the ZymoBIOMICS RNA Miniprep kit (Zymo Research) with 40 s of bead beating at 6 m/s. The RNA samples were pure as indicated by ratios Abs_260_/Abs_280_ ∼ 2.0 and Abs_260_/Abs_230_ > 2.1. Electrophoreses on agarose gel showed no RNA degradation and complete DNA digestion. 600 ng of RNA were then retrotranscribed with the ZymoScript RT PreMix kit (Zymo Research). The extension was performed at 50 °C for 15 min to facilitate the synthesis of high-GC cDNAs. qPCRs (two technical replicates) were performed with the Fast SYBR Green Master mix (Applied Biosystems) with an optimized pair of primers for *OVOA1* and using the endogenous control *CβLP* (Cre06.g278222_4532.1; Dannay et al. 2025). Primers sequences are provided in Supplementary Table S7. Expression fold changes were calculated with the 2^−ΔΔCт^ method (Livak and Schmittgen 2001).

### Quantification of photosynthetic activity

The *ovoa1* mutants and reference strains were grown in TP medium at 20 °C under continuous light (50 µmol photons m^-2^ s^-1^) and stirring (180 rpm) for 4 days. After dilution to 1 x 10^6^ cells/mL with spent TP medium, the cultures were treated with 6 µM Rose Bengal in low light (50 µmol photons m^-2^ s^-1^) for 6 h. The maximum quantum efficiency of photosystem II (F_v_/F_m_) was quantified using a Handy PEA+ chlorophyll fluorometer (Hansatech Instruments). After dark-adapting the cells for 5 min, a light pulse of 3500 μmol photons m^-2^ s^-1^ with an auto-gain was applied for 10 s.

### Multiple sequence alignment

*C. reinhardtii* OVOA1 (Cre08.g380000_4532.1.p) and other non-heme iron sulfoxide/selenoxide synthases protein sequences (from Kayrouz et al. 2024 and references cited within) were aligned using Muscle5 5.3 (Edgar 2022). The alignment was manually edited with SnapGene (www.snapgene.com) and functional residues were annotated based on Kayrouz et al. 2024 and Ireland et al. 2025 using ggmsa (Zhou et al. 2022). The sulfoxide synthase and methyltransferase domains were annotated with interproscan 5.71-102.0 (Blum et al. 2021).

### *OVOA1* homolog occurrence

Archaeplastida proteomes were downloaded from the sources referenced in Supplementary Table S1. When multiple proteomes from the same species were available, they were concatenated. Homologs of *C. reinhardtii* OVOA1 (Cre08.g380000_4532.1.p) were retrieved using diamond 2.1.10 (Buchfink et al. 2021) in ultra-sensitive mode. Taxonomy was assigned with taxize (Chamberlain and Szöcs 2013) using the NCBI taxonomy database. BUSCO analysis of each proteome was performed using compleasm 0.2.7 (Huang and Li 2023) with the OrthoDB 12 database for eukaryotes (Tegenfeldt et al. 2025). The species tree was adapted from Cheng et al. 2019; Gawryluk et al. 2019; Li et al. 2020; Schön et al. 2021.

### OVOA1 phylogeny

The reconstruction of the phylogenetic tree of OvoA/OVOA1 homologs was done with a Snakemake 9.11.2 workflow (Mölder et al. 2021) similar to TreeTuner (Zhang et al. 2022). Sequence manipulations were done in R language or with SeqKit2 (Shen et al. 2024). The OVOA1 5-histidylcysteine sulfoxide synthase domain (NCBIFAM: TIGR04344; InterPro: IPR027577) was extracted from *C. reinhardtii* OVOA1 (Cre08.g380000_4532.1.p) with interproscan 5.74-105.0 (Blum et al. 2021). This domain sequence was used to search for homologs in a database for Archaeplastida (Supplementary Table S1) that we build and the UniProtKB/TrEMBL database with diamond 2.1.11 (Buchfink et al. 2021). Only the aligned part of each hit (corresponding to the 5-histidylcysteine sulfoxide synthase domain) was retrieved due to the variability of the OvoA domain architecture, *e.g.* fusion with OvoB (Gerdol et al. 2019) or missing methyltransferase domain (Ireland et al. 2025). First, a large unrooted approximate tree was built with all these sequences, which were aligned together with FAMSA 2.4.1 (Gudyś et al. 2025). The multiple sequence alignment was trimmed with ClipKIT 2.4.1 (Steenwyk et al. 2020) in gappy mode (threshold 0.9) and compositional bias was reduced with WitChi (commit 3094f08 of https://github.com/stephkoest/witchi; Köstlbacher et al. 2025). The WAG substitution model was determined to be best by ModelFinder (Kalyaanamoorthy et al. 2017) and used to build an approximately maximum-likelihood tree with FastTree 2.1.11 (Price et al. 2010). Long branches and redundant bacterial branches were pruned with TreeShrink 1.3.9 (Mai and Mirarab 2018) and Treemmer 0.3 (Menardo et al. 2018), respectively, to reach a final number of 2090 sequences. The subset of sequences was used to reconstitute a second precise tree. The multiple sequence alignment was built and trimmed the same way as before except that the gap threshold was increased to 0.98. Compositional bias was also reduced with WitChi, allowing a maximum of 10% of the columns to be removed. The final maximum-likelihood tree was built with IQ-TREE 2.4.0 (Minh et al. 2020) with 1000 UFBoot2 replicates (Hoang et al. 2018) and the Q.pfam+I+R10 model (Minh et al. 2021) which was determined to be the best among nuclear protein models by ModelFinder (Kalyaanamoorthy et al. 2017). Some ergothioneine synthase sequences were also recovered in this analysis and, as expected (Kayrouz et al. 2024), were placed as outgroup by minimal ancestor deviation / minimum variance rooting with mad 2.2 (Tria et al. 2017; Mai et al. 2017). This outgroup clade was collapsed and the tree was visualized with tidytree, ggtree, and treeio (Wang et al. 2020).

### Microarrays and RNA-seq analysis

*C. reinhardtii* probes (Gene Expression Omnibus accession: GPL13913; Fischer et al. 2012) were remapped to the version 6.1 of the transcriptome using minimap2 2.28 (Li 2018) with the “sr” preset. The package limma 3.64.3 (Phipson et al. 2016) was used for background-correction and quantile-normalization of microarray data (Gene Expression Omnibus accession: GSE30648; Fischer et al. 2012), as well as differential expression analysis.

RNA-seq of the 4A wild type and the *hy5-3*, *co-2*, and *hy5-co* mutants was performed after 0, 15, 30, and 60 min of high light (350 µmol photons m^-2^ s^-1^) in HSM medium (Hui et al. 2023). For *spa1*, RNA-seq was performed with cells grown in HSM in low light (60 µmol photons m^-2^ s^-1^). These experiments are detailed in another publication (Benko et al.). Newly generated or publicly available RNA-seq datasets were processed with a Snakemake 9.11.2 workflow (Mölder et al. 2021). Briefly, published reads were retrieved from NBCI SRA with kingfisher 0.4.1 (Woodcroft et al. 2024) or directly from the Genome Sequence Archive (https://ngdc.cncb.ac.cn/gsa/). Preprocessing and quality control was done with fastp 0.24 (Chen 2023) and the transcripts abundance was quantified with Salmon 1.10.3 (Patro et al. 2017) against the version 6.1 of the *C. reinhardtii* genome annotation (Craig et al. 2023). MultiQC 1.31 was used for quality control of preprocessing and quantification (Ewels et al. 2016). DESeq2 1.44 was used for count normalization and differential expression analysis (Love et al. 2014). In each dataset, the condition/timepoint where *OVOA1* (Cre08.g380000_4532.1) was expressed the most in the wild type was chosen to calculate the fold changes and false discovery rates reported in Figure 7D.

For the co-expression analysis, TPM (transcripts per million) obtained with Salmon were mean-centering normalized as in Salomé and Merchant 2021 with preprocessCore 1.66 (Bolstad 2025) before calculating Pearson correlation coefficients. Protein subcellular localization was predicted with DeepLoc 2.1 (Ødum et al. 2024) and their classification was done manually with information from *C. reinhardtii* genome annotation (Craig et al. 2023), annotation done with eggNOG-mapper 2.1.13 (Cantalapiedra et al. 2021), ARC mutants (Wakao et al. 2021), and *A. thaliana* (Araport11; Reiser et al. 2024) homologs recovered with diamond 2.1.11 (Buchfink et al. 2021). Sequence manipulations and annotation version reconciliation were performed using R language, MMseqs2 (Steinegger and Söding 2017), and SeqKit2 (Shen et al. 2024).

## Acknowledgements

We thank Natalie Heimerl for backcrossing the *ovoa1-1* mutant, Andrei Ogonkov for providing us with a protocol for ovothiol A extraction, Thomas Roach for early access to RNA-seq data, Sunnyjoy Dupuis and Donat Wulf for their advice on RNA extraction and bioinformatics analyses, and Zhongrui Zhou and Andrea Perner for performing part of the LC-MS analyses depicted in Figure 6B and Supplementary Figure S6B, respectively. We are grateful to Vincent Boudreau, Derrick S. Chuang, Marilyn Kobayashi, Christophe H. Marchand, Maria Mittag, Claire Remacle, Melissa S. Roth, Cailyn Sakurai, Srikanth Tirumani, Shivani Upadhyaya, and William Zerges for stimulating discussions and support.

## Author contributions

Y.L., F.C., F.P.S., K.K.N., S.W., and S.S. designed experiments; Y.L., F.C., G.S.A, E.H., M.H., G.B., S.C.S., and S.S. performed experiments; R.N., M.G., C.H., and A.R.G. contributed methods; Y.L., F.C., S.W., and S.S. wrote the manuscript; all authors analyzed data, edited the manuscript, and approved its final version.

## Supplementary data

The following materials are available in the online version of this article.

**Supplementary Table S1.** Archaeplastida genomes and transcriptomes used for the phylogenetic analysis of OvoA/OVOA1.

**Supplementary Table S2.** Strains used in this study.

**Supplementary Table S3.** Viability of the *ovoa1-4* mutant in response to ROS-inducing treatments.

**Supplementary Table S4.** Description of the top 150 genes co-expressed with *OVOA1*.

**Supplementary Table S5.** RNA-seq metadata and *OVOA1* fold change and adjusted *p*-value in mutants compared to reference strains.

**Supplementary Table S6.** Sequences of crRNAs and single-stranded DNA templates (ssODNs) for CRISPR-Cas9 mutagenesis.

**Supplementary Table S7.** Primers used in this study.

**Supplementary Table S8.** Occurrence of OvoA/OVOA1 orthologs in Archaeplastida.

**Supplementary Table S9.** F_v_/F_m_ of the *ovoa1-4* mutant in response to Rose Bengal.

**Supplementary Table S10.** Expression of *OVOA1* in response to high light.

**Supplementary Table S11.** Expression of *OVOA1* in response to Rose Bengal.

**Supplementary Table S12.** Relative ion counts of Ovothiol A during singlet oxygen acclimation.

**Supplementary Table S13.** Expression of *OVOA1* during singlet oxygen acclimation.

**Supplementary Table S14.** Predicted localization of the top 150 *OVOA1*-co-expressed genes.

**Supplementary Table S15.** DEseq2 normalized counts of *OVOA1* during high light treatment.

**Supplementary Figure S1.** Derivatization reactions of ovothiol A.

**Supplementary Figure S2.** Quantification of ovothiol A.

**Supplementary Figure S3.** Chlamydomonas OVOA1 is a functional 5-histidylcysteine sulfoxide synthase.

**Supplementary Figure S4.** Screening for CRISPR-mediated *ovoa1* mutants by colony PCR and restriction analysis.

**Supplementary Figure S5.** Sanger sequencing of CRISPR-mediated *ovoa1* mutants.

**Supplementary Figure S6.** *ovoa1* insertional mutants are impaired in ovothiol A production.

**Supplementary Figure S7.** *OVOA1* basal transcript level under standard low light conditions.

**Supplementary Figure S8.** Growth curves of *ovoa1* mutants and reference strains.

**Supplementary Figure S9.** *OVOA1* is induced by light and potentially by the circadian cycle during the diurnal cycle.

**Supplementary Figure S10.** Growth of *ovoa1* mutants and reference strains on Rose Bengal-containing plates.

**Supplementary Figure S11.** The *ovoa1-1* and *ovoa1-2* mutants are sensitive to singlet oxygen.

**Supplementary Figure S12.** Identification of genes co-expressed with *OVOA1* in a transcriptome-wide matrix.

**Supplementary Figure S13.** Analysis of the genes co-expressed with *OVOA1*.

**Supplementary Dataset S1.** *OVOA1* and *ALS1* allele sequences for the *ovoa1* mutants and reference strains generated by CRISPR-Cas9 editing.

**Supplementary Dataset S2.** Sequence of His-tagged OVOA1 used for recombinant protein production.

## Funding

Funding by the German Research Foundation (DFG; grant no. SA 2453/3-1 to S.S. and project ID 239748522 - SFB 1127 to C.H. and S.S.), the Rose Hills Foundation (to S.C.S.), the Howard Hughes Medical Institute (to K.K.N.), and the Swiss National Science Foundation (grants no. 51NF40-205608 and 219624 to F.P.S.) is gratefully acknowledged. S.W. was supported by the U.S. Department of Energy, Office of Science, through the Photosynthetic Systems program in the Office of Basic Energy Sciences. Y.L. was supported by a personal Ph.D. fellowship from Leipzig University (*Doktorandenförderplatz*). The LC-MS device was funded by the DFG (INST 268/451-1 FUGG) and the Saxon Ministry for Science, Culture and Tourism (SMWK).

## Conflict of interest statement

The authors declare that they have no conflict of interest.

## Data availability

Additional data and scripts used in the transcriptomics and phylogenetic analyses are available at the following link: https://doi.org/10.5061/dryad.2bvq83c4x.

## Notes

https://doi.org/10.5061/dryad.2bvq83c4x

### Competing Interest Statement

The authors have declared no competing interest.

### Summary of Updates

The ORCID of one of the author was referring to another researcher with the same name so it was corrected.

