## Supplementary Figures and Datasets for "Ovothiol A mediates singlet oxygen resistance and acclimation in Chlamydomonas"

### Table of Contents

Supplementary Figures	2

Supplementary Datasets	16

Supplementary References	19

### Supplementary Figures

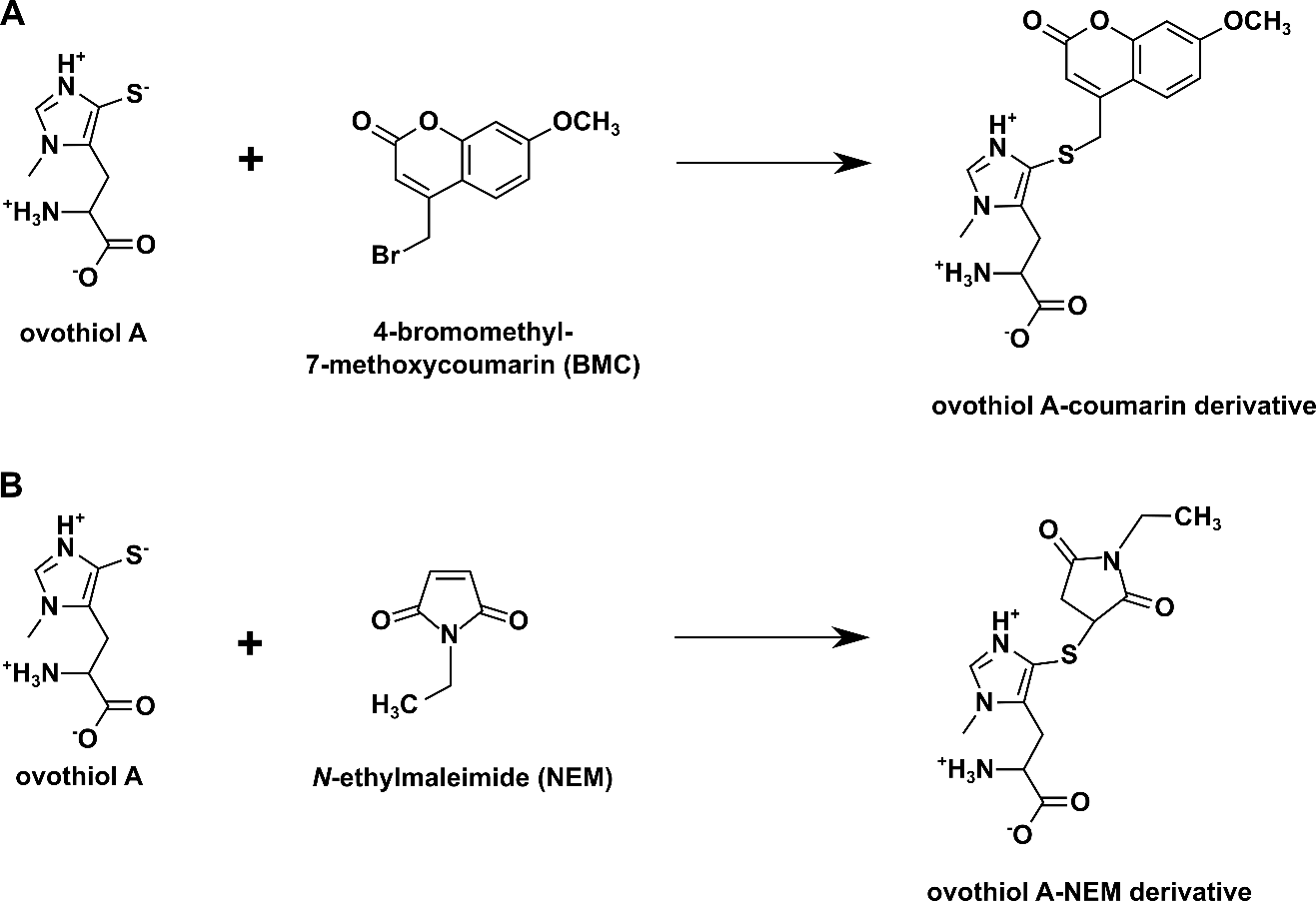

#### Supplementary Figure S1: Derivatization reactions of ovothiol A.

(A) Derivatization of ovothiol A with 4-bromomethyl-7-methoxycoumarin (BMC). Reduced ovothiol A reacts with BMC via nucleophilic substitution. (B) Derivatization of ovothiol A with *N*-ethylmaleimide (NEM) to distinguish between the reduced and oxidized ovothiol A. Reduced ovothiol A reacts with NEM via a Michael addition. This derivatization was used to prevent the autoxidation of reduced ovothiol A and detect the underivatized oxidized ovothiol A using an NH_2_ column.
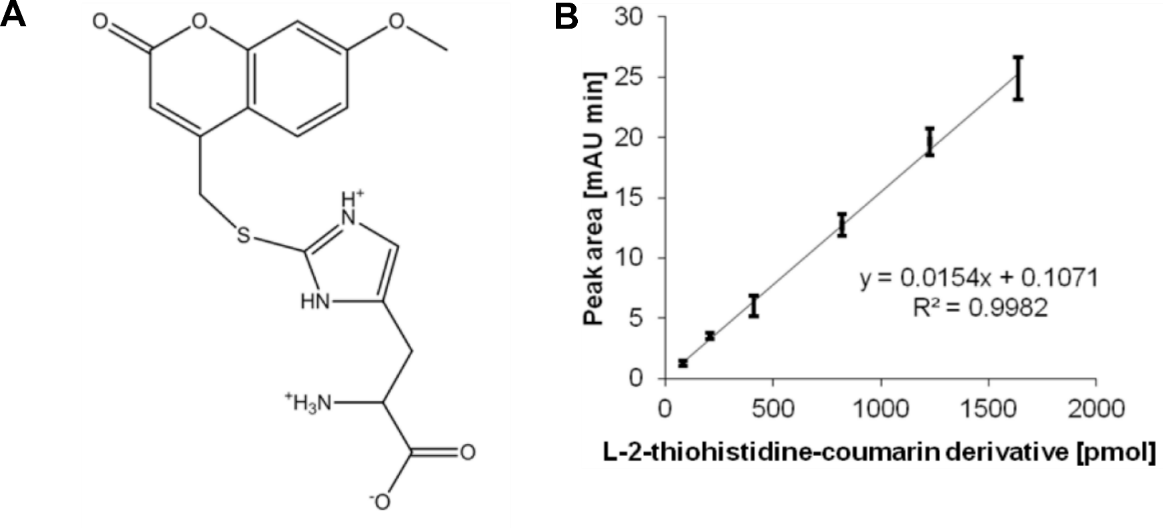

#### Supplementary Figure S2: Quantification of ovothiol A.

(A) Structural formula of L-2-thiohistidine-coumarin derivative. L-2-thiohistidine was derivatized with BMC in the same way as ovothiol A. Because of the addition of the UV light-absorbing methoxycoumarin moiety, the derivatization product can be detected by HPLC with an absorption detector at 330 nm. (B) Standard curve of L-2-thiohistidine-coumarin used for the quantification of intracellular ovothiol A. Mean ± standard deviation of three replicates is shown.

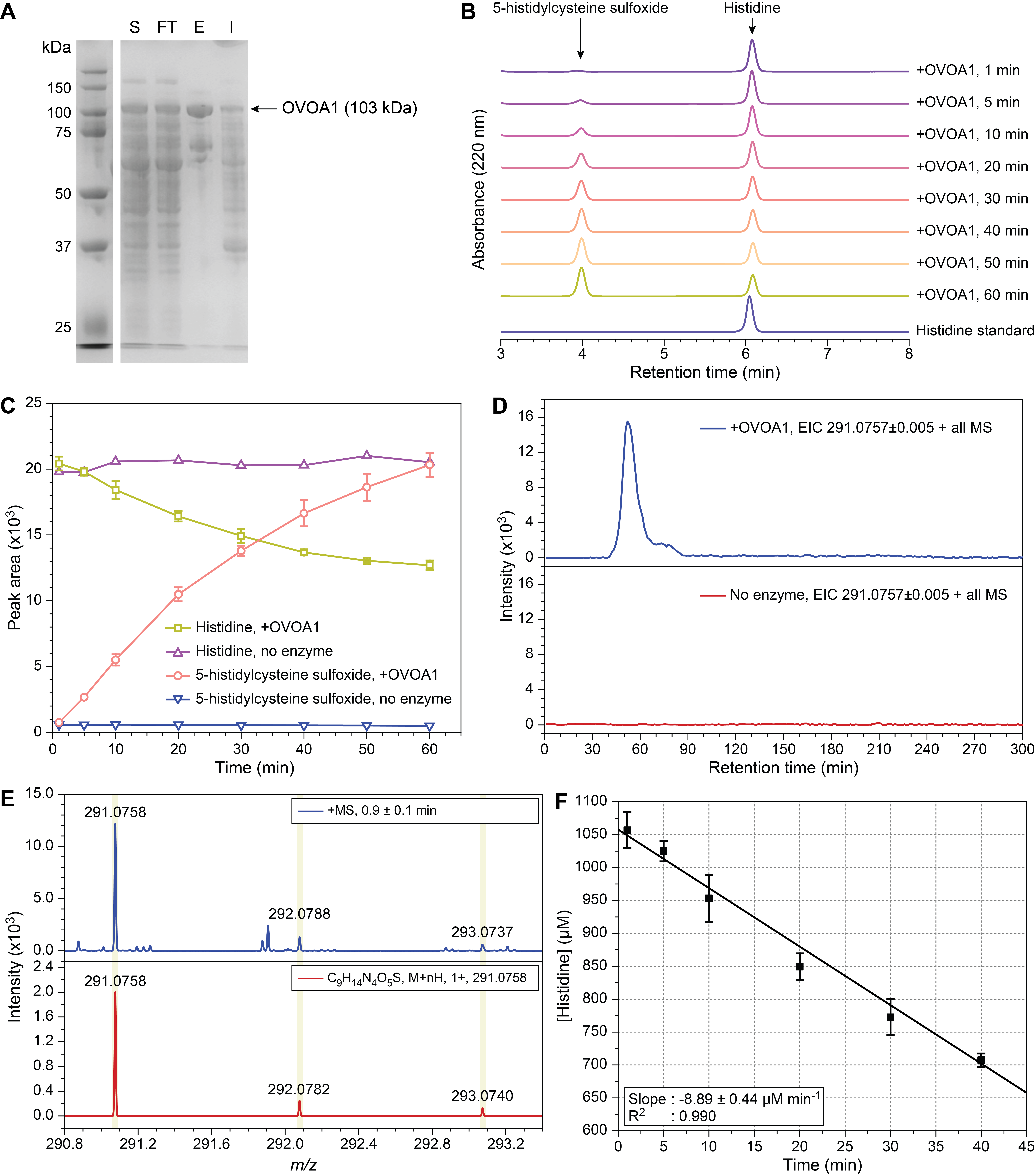

#### Supplementary Figure S3: Chlamydomonas OVOA1 is a functional 5-histidylcysteine sulfoxide synthase.

(A) SDS-PAGE of His-tagged Chlamydomonas OVOA1 recombinantly produced in *E. coli* and purified on a Ni-NTA column. S: cell supernatant after centrifugation; FT: flowthrough; E: eluted fraction; I: insoluble fraction suspended in urea. *(legend continued on next page)*

(B) HPLC chromatograms of OVOA1 reaction with cysteine (not shown) and histidine over 60 min, and (C) peak area of the substrate L-histidine (abbreviated as “histidine”) and the product 5-L-histidyl-L-cysteine sulfoxide (abbreviated as “5-histidylcysteine sulfoxide”). See Figure 1B for the reaction schematic and chemical structures. (D) The formation of 5-histidylcysteine sulfoxide (*m/z* calc. = 291.0758; *m/z* obs. = 291.0758) was confirmed by LC-MS. The extracted ion chromatograms (EIC) of reactions with (blue) and without (red) OVOA1 shows that 5-histidylcysteine sulfoxide (*m/z* calc. = 291.0758) is formed only in the enzyme-containing reaction. (E) The observed isotope pattern of 5-histidylcysteine sulfoxide matches the predicted pattern. (F) Determination of the observed rate of OVOA1-catalyzed consumption of histidine. OVOA1 concentration was 1.0 μM, the product formation was estimated at 8.890 ± 0.44 μM min^-1^, the calculated initial rate was 0.148 ± 0.007 s^-1^, and the calculated specific activity was 0.086 ± 0.004 μmol min^-1^ mg^-1^. For (C) and (F), data points represent mean ± standard deviation of triplicates.

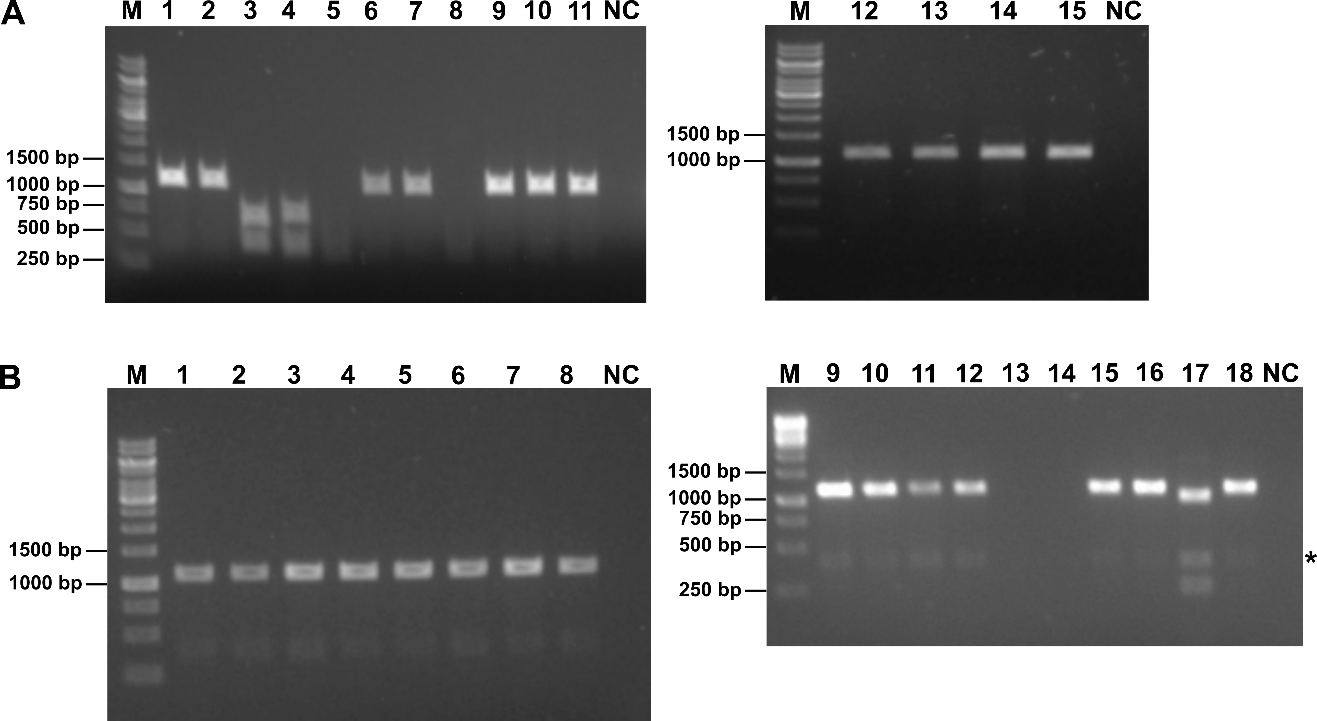

#### Supplementary Figure S4: Screening for CRISPR-mediated *ovoa1* mutants by colony PCR and restriction analysis.

(A) Identification of *ovoa1* mutants edited in exon 2. A fragment of the *OVOA1* gene (Cre08.g380000_4532.1) with an expected size of 1070 bp was amplified by colony PCR of 15 transformants. The PCR products were digested with *Eco*RV (lanes 1-15). The expected sizes of the digestion products were 650 bp and 420 bp. Sequencing of the PCR products confirmed the premature stop codons in clones 3 and 4, which correspond to *als1-2 ovoa1-4* (strain SSJL-1) and *als1-1 ovoa1‑5* (strain SSJL-2) (Dataset S1). Clone 4 (*als1-1 ovoa1‑5*) had an additional insertion and duplication resulting in three *Eco*RV restriction sites; therefore, one band after restriction digestion ran higher in the gel than expected (702 bp instead of 650 bp based on the sequencing results), whereas the remaining digestion products of 73 bp and 90 bp were not visible in the 1% agarose gel. (B) Identification of *ovoa1* mutants edited in exon 3. A fragment of the *OVOA1* gene with an expected size of 1070 bp was amplified by colony PCR of 18 transformants. PCR products were then digested with *Nhe*I (lanes 1-18). The expected sizes of the digestion products were 832 bp and 238 bp. Sequencing of the PCR product confirmed the premature stop codon in clone 17, which corresponds to *als1-1 ovoa1-6* (strain SSJL-3) (Dataset S1). Clone 17 (*als1-1 ovoa1-6*) had an additional insertion and duplication resulting in two *Nhe*I restriction sites; therefore, one band after restriction digestion ran higher in the gel than expected (947 bp instead of 832 bp based on the sequencing results), whereas the remaining digestion product of 48 bp was not visible in the 1% agarose gel. See Supplementary Table S7 for primer sequences. NC, negative control with water instead of DNA; M, GeneRuler 1 kb DNA Ladder; *, unspecific PCR product.

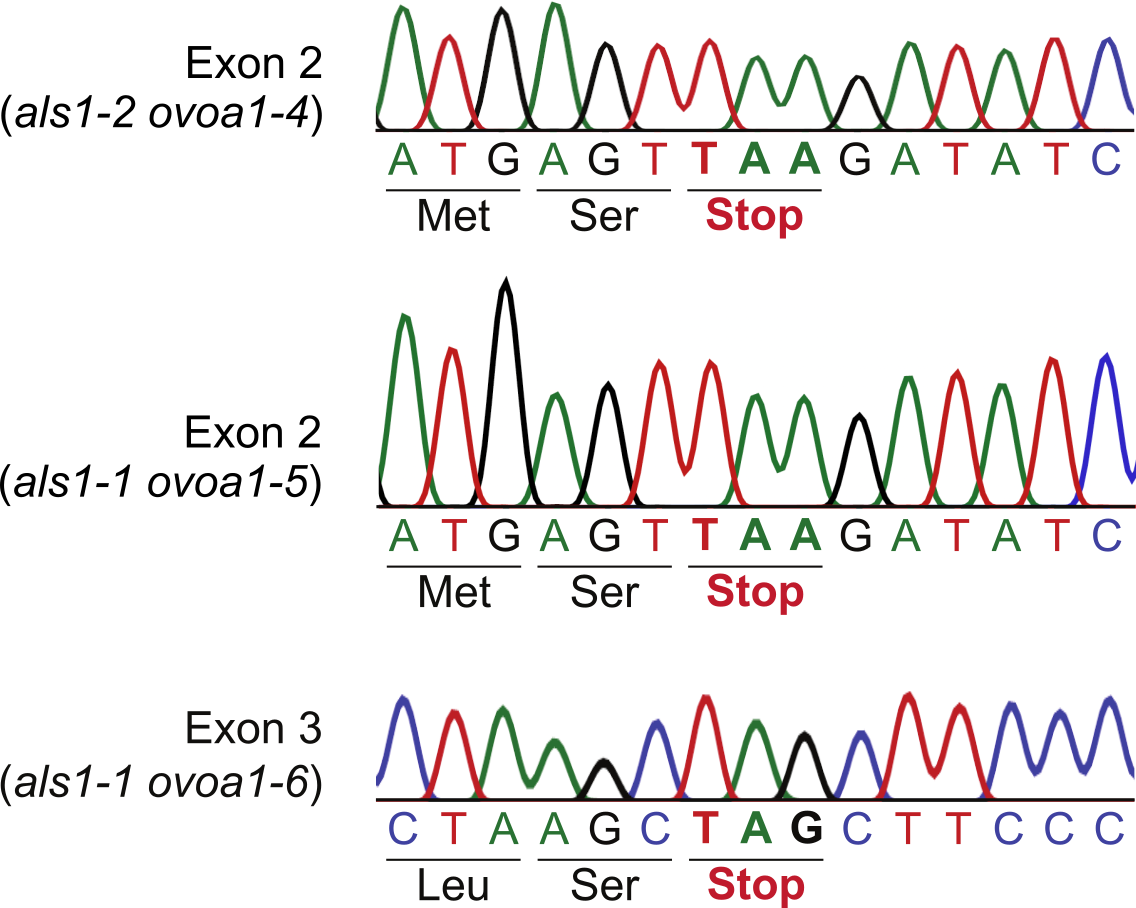

#### Supplementary Figure S5: Sanger sequencing of CRISPR-mediated *ovoa1* mutants.

The relevant sections of the electropherograms containing the premature stop codons in exons 2 or 3 of *OVOA1* are shown.

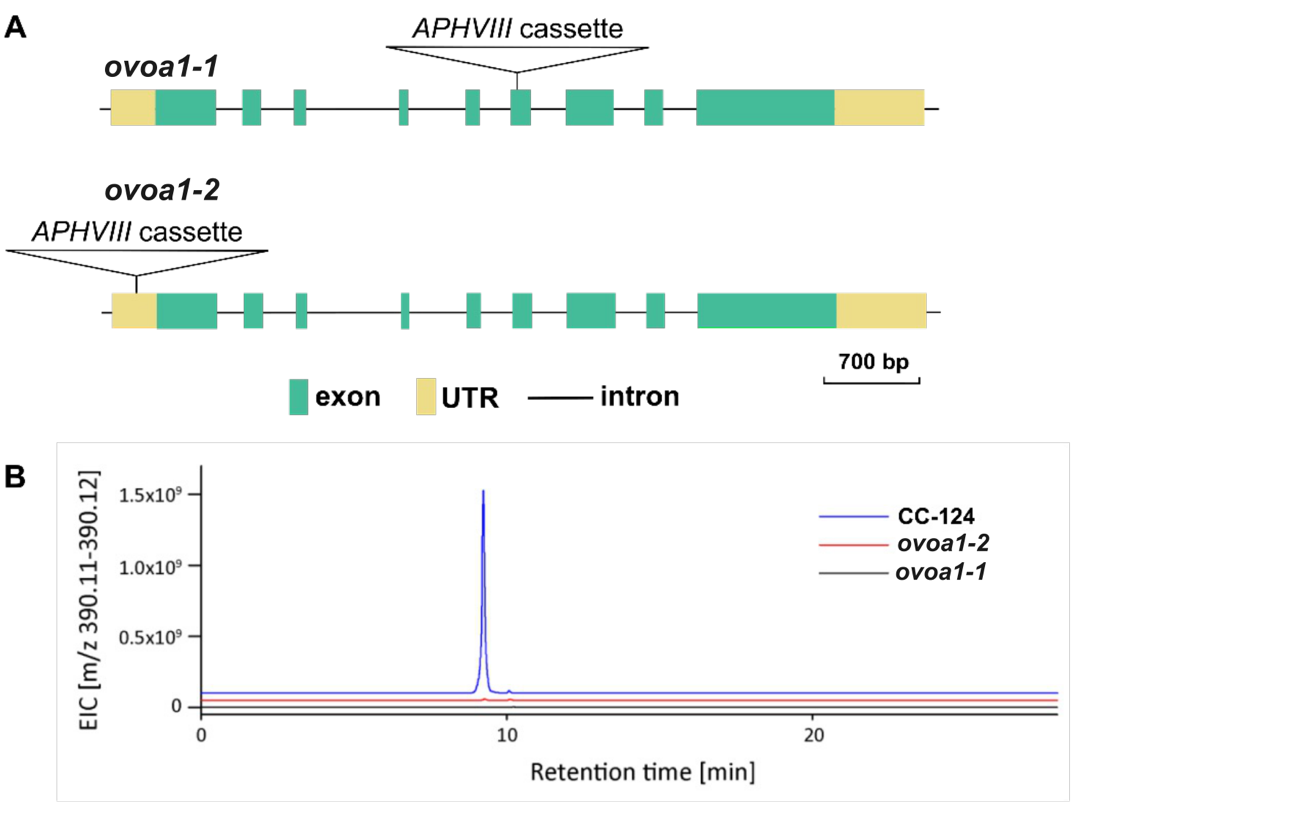

#### Supplementary Figure S6: *ovoa1* insertional mutants are impaired in ovothiol A production.

(A) Genotypes of the *ovoa1* insertional mutants. The *ovoa1-1* mutant was isolated by PCR screening of around 25,000 random-insertion mutants and contains an insertion of the *APHVIII* cassette in the 6^th^ exon of the *OVOA1* gene (Cre08.g380000_4532.1). The *ovoa1-2* mutant was obtained from the CLiP library (LMJ.RY0402.159407), backcrossed with CC-124 and CC-125, and contains an insertion of the *APHVIII* cassette in the 5'-untranslated region (UTR). (B) Extracted ion chromatograms (EIC) for coumarin derivative of ovothiol A in the wild type CC-124 and the *ovoa1* insertional mutants. In CC‑124, ovothiol A-coumarin was detected (observed: *m/z* 390.11155; expected for protonated ovothiol A-coumarin C_18_H_20_O_5_N_3_S^+^: *m/z* 390.11182, deviation 0.69 ppm). This peak is absent from *ovoa1‑1*, while a very small peak is visible in *ovoa1‑2* with a peak area of less than 1% compared to the wild type (observed: *m/z* 390.11137, deviation 1.2 ppm).
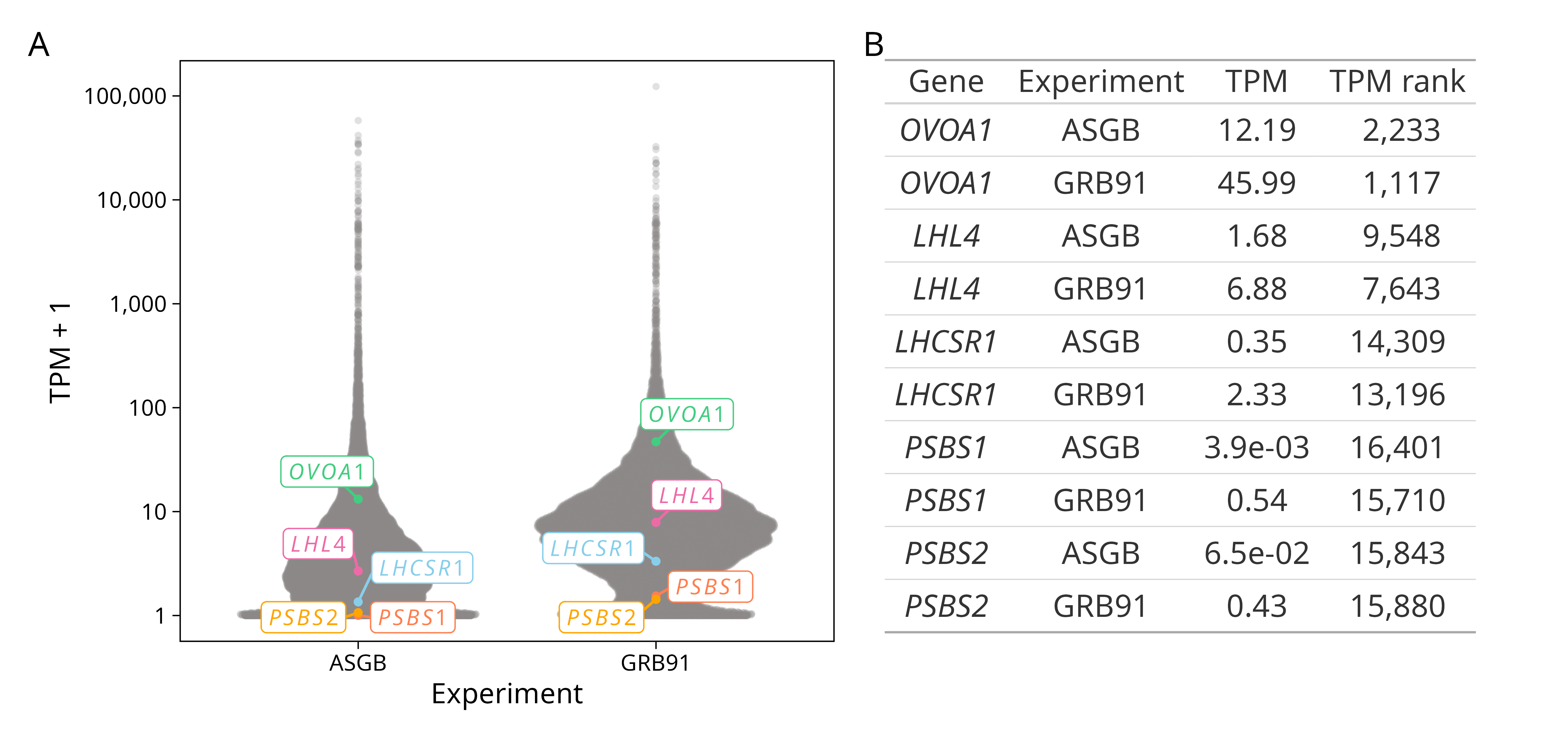

#### Supplementary Figure S7: *OVOA1* basal transcript level under standard low light conditions.

TPM (transcripts per million) were quantified in two independent RNA-seq experiments (ASGB and GRB91) done with the 4A wild type grown autotrophically (HSM medium) in low light (60 µmol photons m^-2^ s^-1^). The data points are averages calculated from at least three biological replicates. (A) TPM+1 of all Chlamydomonas genes (in gray) with *OVOA1* and other CONSTANS-dependent photoprotection genes highlighted (Gabilly et al. 2019; Dannay et al. 2025). (B) TPM and TPM rank values of selected genes. TPM ranks are the ranks of all 17,616 genes when ordered from high to low TPM. With these values, *OVOA1* transcript ranked in the top 6.3-12.7% of the most highly expressed genes. *LHH4*, Light Harvesting complex-Like 4 (Cre17.g740950_4532.1); *LHCSR1*, Light-Harvesting Complex Stress-Related 1 (Cre08.g365900_4532.1); *PSBS1*, Photosystem II-associated 22 kDa protein S 1 (Cre01.g016600_4532.1); *PSBS2*, Photosystem II-associated 22 kDa protein S 2 (Cre01.g016750_4532.1).

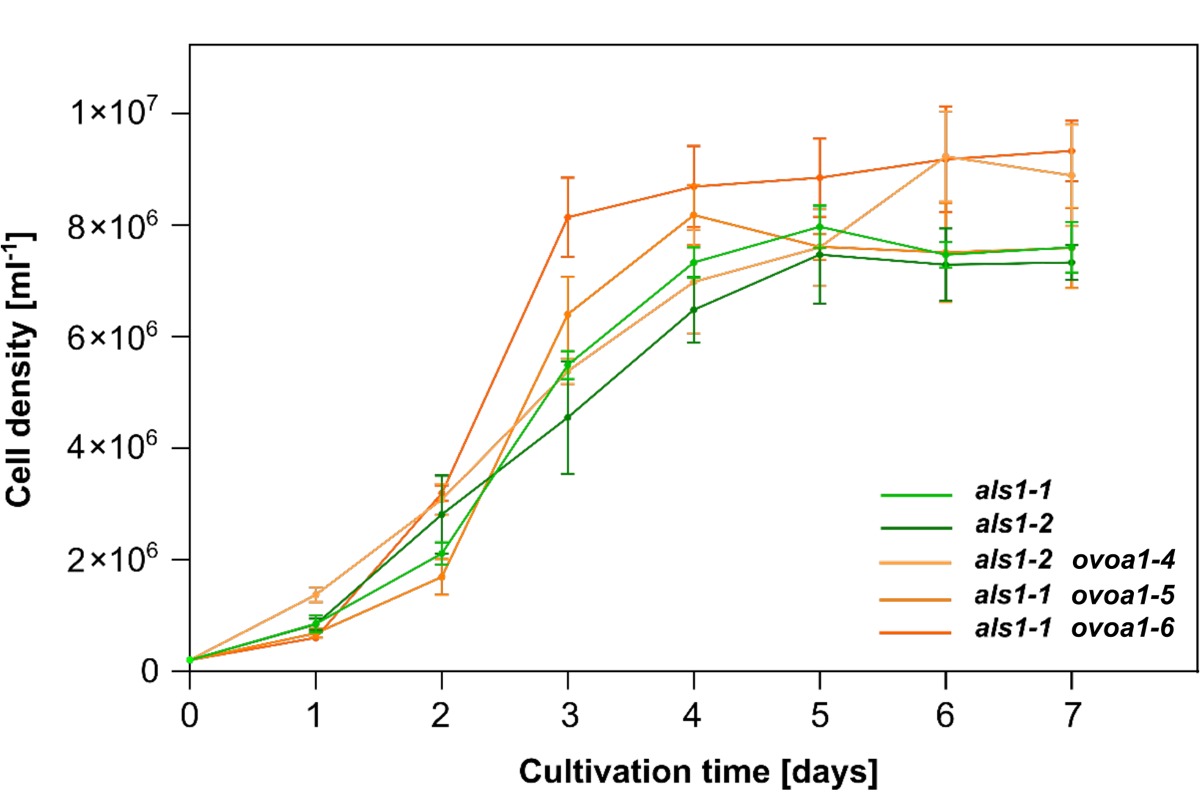

#### Supplementary Figure S8: Growth curves of *ovoa1* mutants and reference strains.

Liquid cultures were incubated at 20 °C under continuous white light (50 µmol photons m^-2^ s^-1^) and stirring (180 rpm) for 7 days. Cell density was measured with a Coulter counter. Values indicate the mean ± standard deviation of three biological replicates.

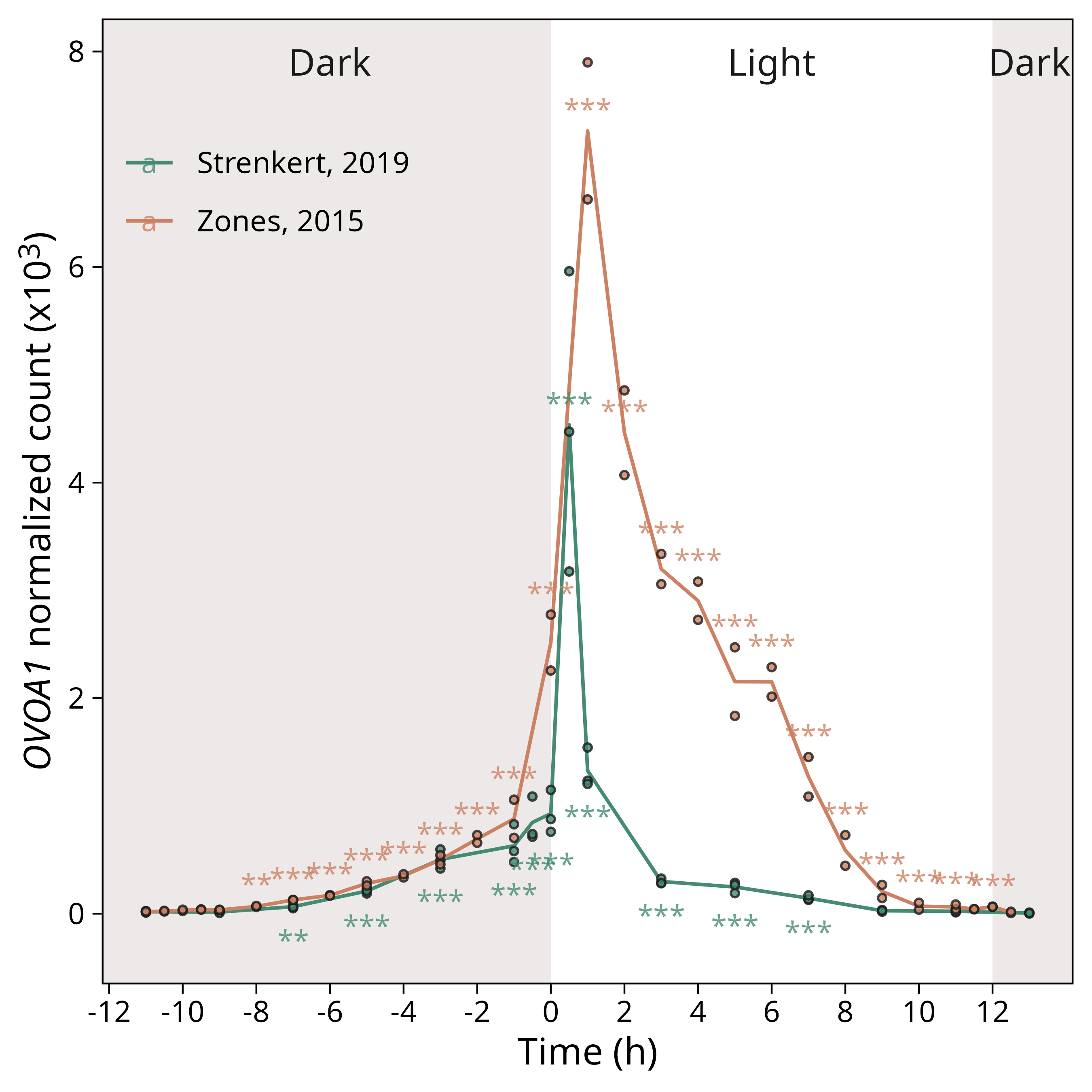

#### Supplementary Figure S9: *OVOA1* is induced by light and potentially by the circadian cycle during the diurnal cycle.

RNA-seq data were re-analyzed and counts were normalized with DESeq2 (Love et al. 2014). The light intensity was 125 µmol photons m^-2^ s^-1^ in Zones et al. 2015 and 200 µmol photons m^-2^ s^-1^ in Strenkert et al. 2019. Circles represent individual biological replicates. Benjamini–Hochberg adjusted *p*-values (false discovery rates) were calculated comparing each timepoint with the -11 h timepoint, and symbols were colored by dataset. * *p* ≤ 0.05; ** *p* ≤ 0.01; *** *p* ≤ 0.001.

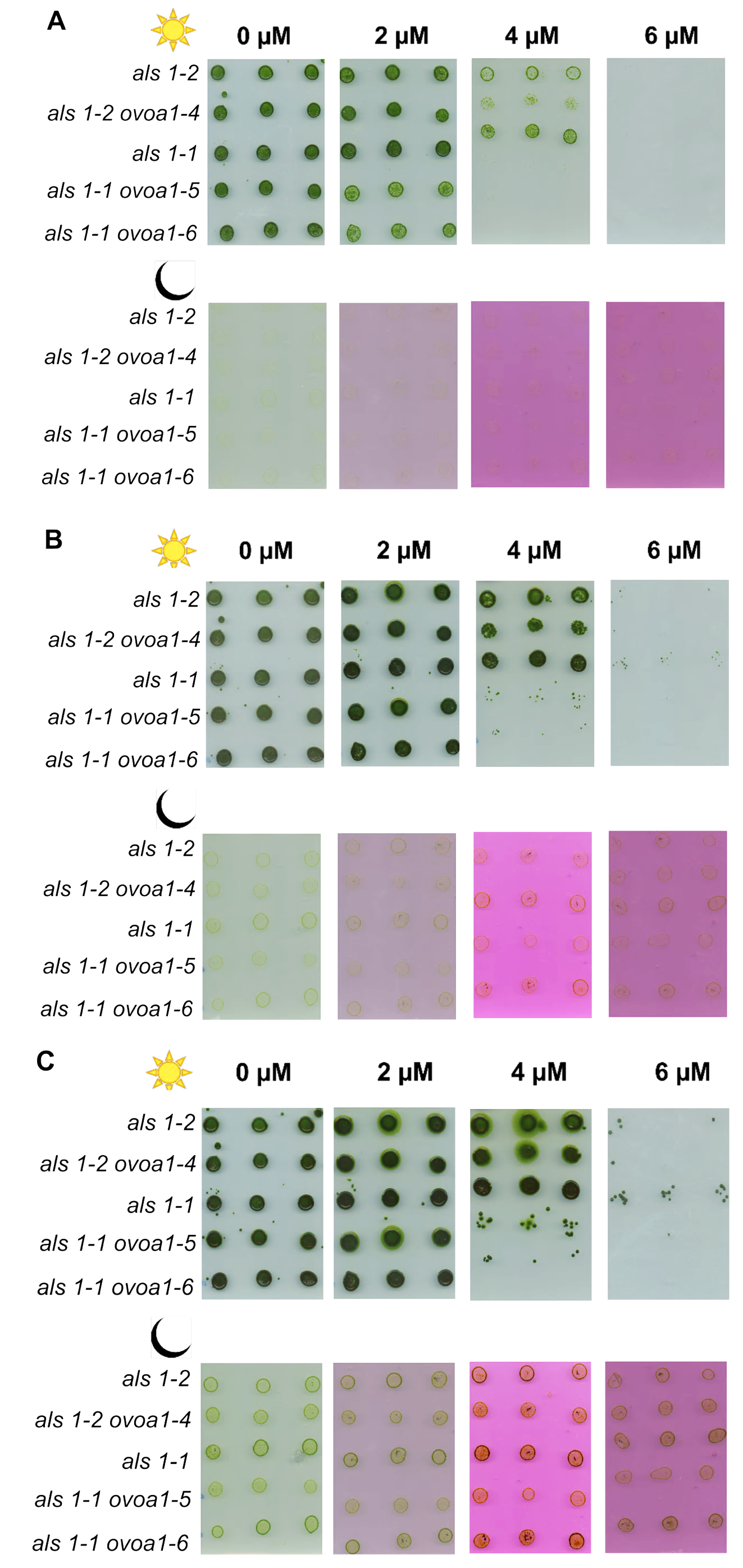

#### Supplementary Figure S10: *legend on the next page.*

#### Supplementary Figure S10: Growth of *ovoa1* mutants and reference strains on Rose Bengal-containing plates.

(A) Sensitivity of *ovoa1* mutants to Rose Bengal after four days. (B) Sensitivity of *ovoa1* mutants to Rose Bengal after six days. (C) Sensitivity of *ovoa1* mutants to Rose Bengal after eight days. TAP plates containing the indicated concentrations of Rose Bengal were inoculated with 2 x 10^4^ cells per spot in three technical replicates. The top rows in (A), (B), and (C) show plates incubated under continuous white light of 50 μmol photons m^-2^ s^-1^ at 20 °C. The bottom rows in (A), (B), and (C) show Rose Bengal-containing control plates incubated in the dark at 20 °C.

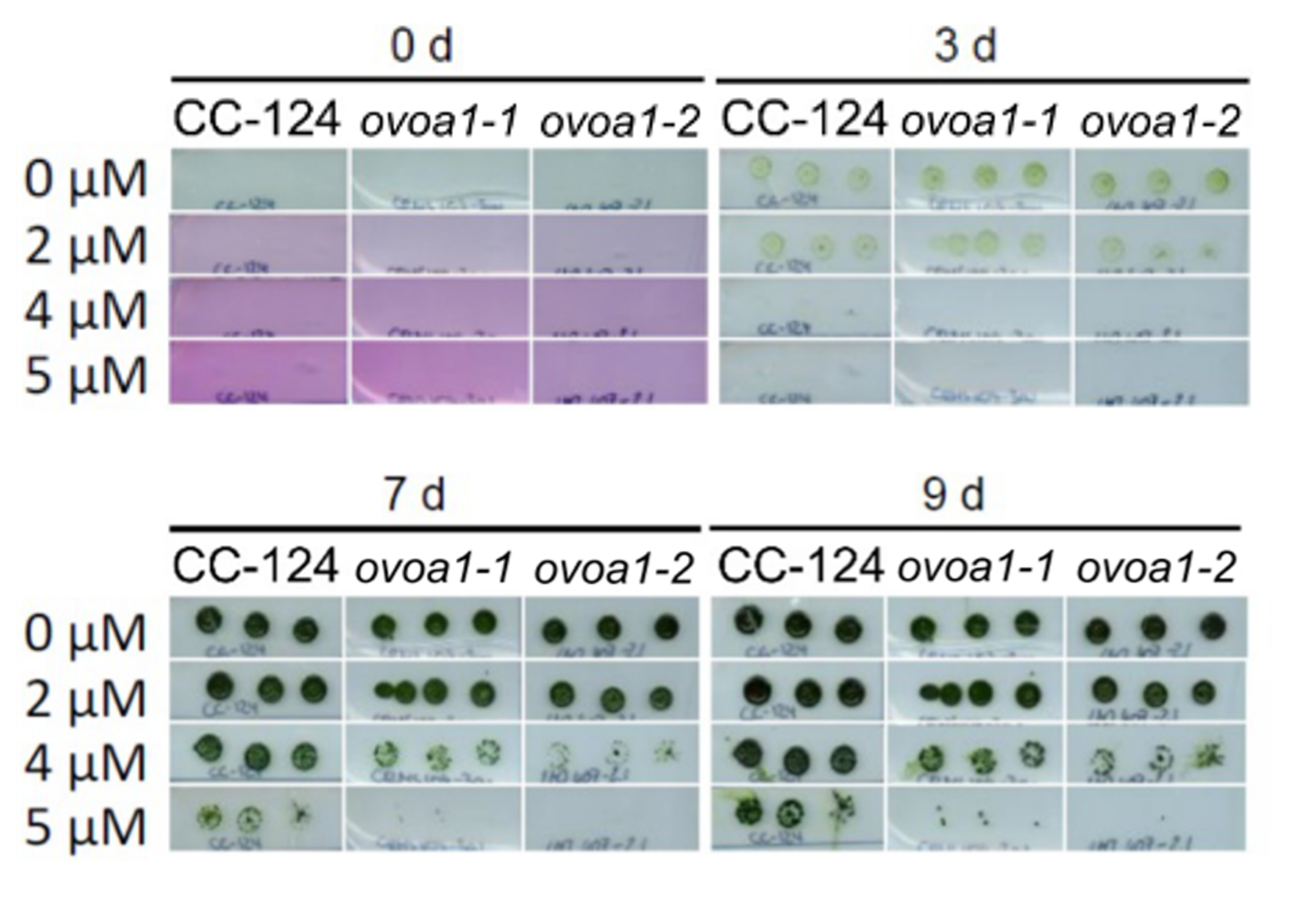

#### Supplementary Figure S11: The *ovoa1-1* and *ovoa1-2* mutants are sensitive to singlet oxygen.

Growth of the wild-type strain CC‑124 and *ovoa1* insertional mutants on Rose Bengal-containing TAP agar plates. TAP plates with the indicated concentrations of Rose Bengal were incubated under continuous white light of 50 μmol photons m^-2^ s^-1^ at 20 °C. For inoculation, 2 x 10^4^ cells were used per spot in three technical replicates.

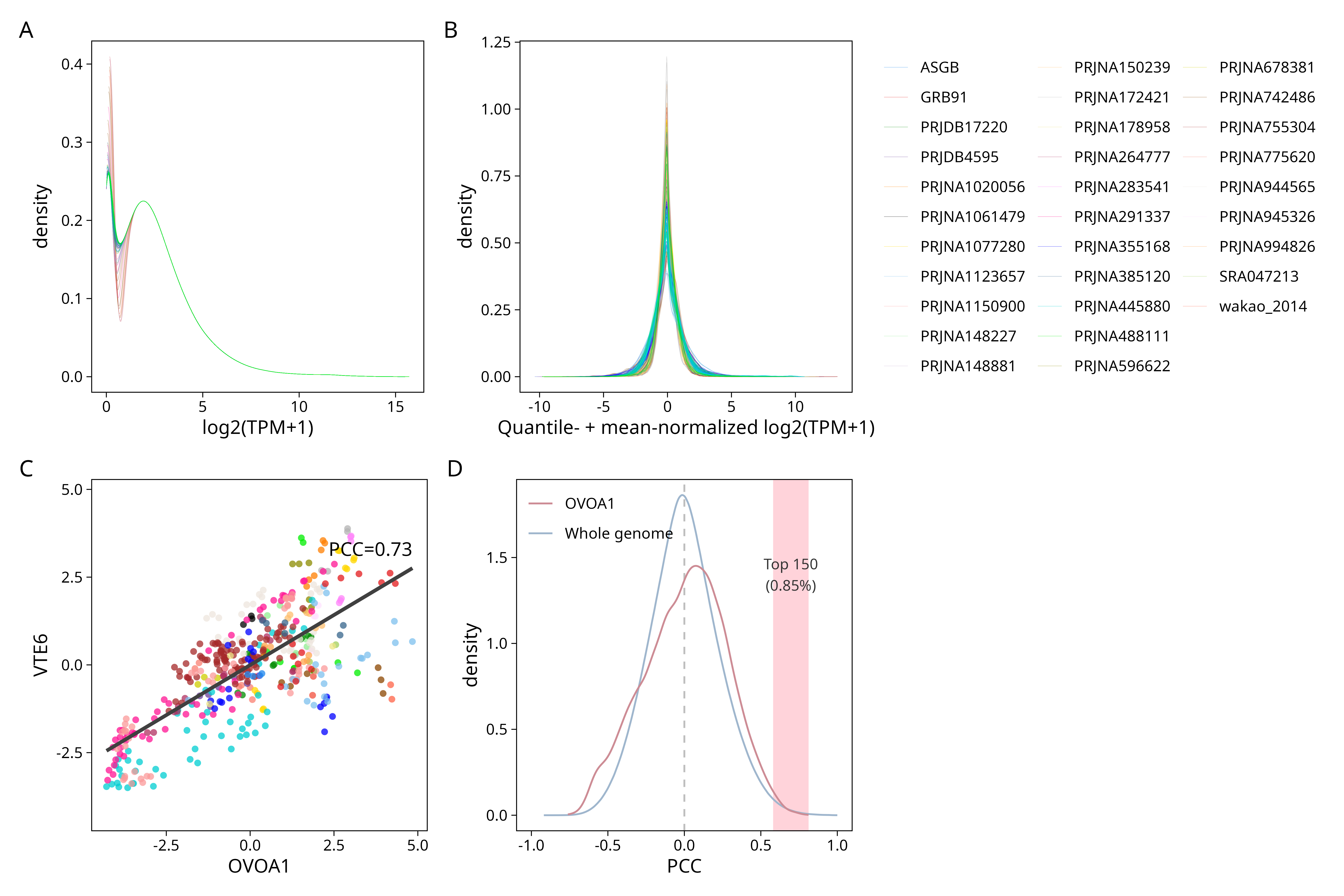

#### Supplementary Figure S12: Identification of genes co-expressed with *OVOA1* in a transcriptome-wide matrix.

461 RNA-seq samples of Chlamydomonas wild types were re-analyzed. TPM+1 (transcripts per million) were log_2_-transformed (A) and quantile- + mean-normalized (B) as in Salomé and Merchant 2021, showing a similar distribution across samples. These fully normalized values were then used to build a co-expression matrix by calculating Pearson correlation coefficients (PCC). (C) Example of a high positive PCC with normalized values for the genes *OVOA1* and *VTE6* (Cre14.g624350_4532.1). Elements in (A), (B), and (C) are colored by dataset. “GRB91” is a newly-generated dataset with the 4A wild type in response to high light (350 µmol photons m^-2^ s^-1^) for 0, 15, 30, and 60 min (Figure 7E; Benko et al. 2025). “ASGB” was generated with the same wild type in low light only (60 µmol photons m^-2^ s^-1^; Benko et al. 2025). “wakao_2014” is the dataset generated with the 4A wild type in Wakao et al. 2014, which is not publicly available. Other datasets are named by their NCBI identifier. (D) Density of PCCs of *OVOA1*-co-expressed genes and all genes annotated in the Chlamydomonas genome. The pink area covers the PCCs of the top 150 genes co-expressed with *OVOA1* which accounts for 0.85% of all genes. Their function is described in detail in Figure 7 and Supplementary Figure S12.

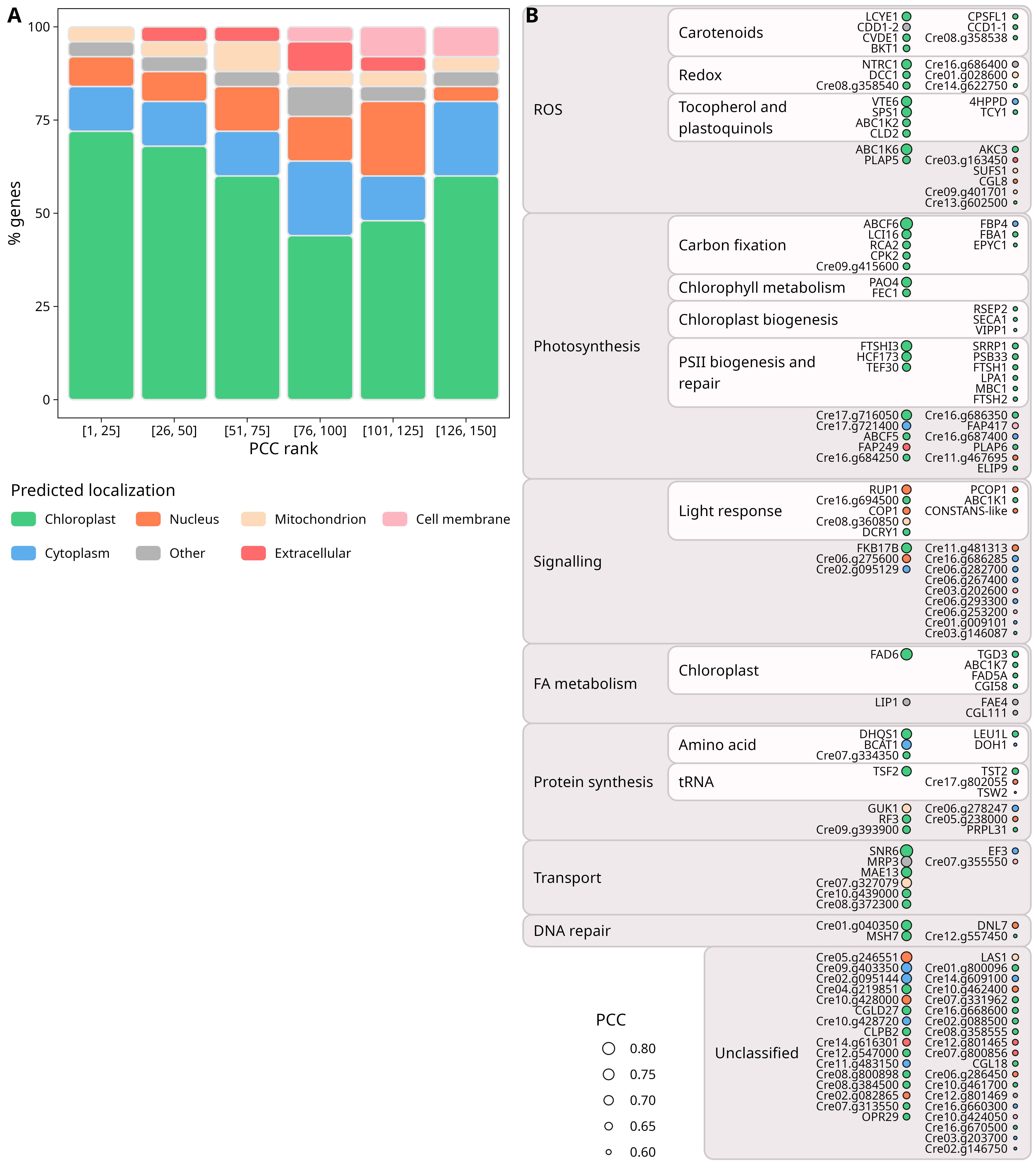

#### Supplementary Figure S13: Analysis of the genes co-expressed with *OVOA1*.

(A) Predicted localization of proteins encoded by *OVOA1* co-expressed genes binned by PCC rank, the smaller the rank the higher the co-expression with *OVOA1*. (B) Classification of the top 150 genes co-expressed with *OVOA1*. See Supplementary Table S4 and Materials and methods for information used to manually annotate the genes. The localization color legend is shared between (A) and (B). “Cre” v6.1 identifiers are abbreviated by removing the strain and transcript numbers. PSII, photosystem II; FA, fatty acid.

### Supplementary Datasets

#### Supplementary Dataset S1: *OVOA1* and *ALS1* allele sequences for the *ovoa1* mutants and reference strains generated by CRISPR-Cas9 editing.

CC-124 was used as a parental strain for the CRISPR/Cas9-mediated co-editing of the *ALS1* (Cre09.g386758_4532.1) and *OVOA1* (Cre08.g380000_4532.1) genes. Reference strains *als1-1 and* *als1-2* were edited only in the *ALS1* gene, whereas strains *als1-2 ovoa1-4*, *als1-1 ovoa1-5*, and *als1-1 ovoa1-6* were edited in both the *ALS1* and *OVOA1* genes. All strains used in this study are summarized in Supplementary Table S2. The *als1-1* allele contains a single nucleotide substitution (AAG -> ACG) in exon 8 of the *ALS1* gene (highlighted in green), causing a missense mutation (K257T). The *als1-2* allele contains two single nucleotide substitutions (AAG -> ACG, highlighted in green; GAC -> GAT, highlighted in magenta); the latter substitution causes a silent mutation (D258) and creates an *Eco*RV restriction site (GATATC, underlined). The *ovoa1-4* allele contains a premature stop codon TAA (highlighted in red) and an *Eco*RV restriction site in exon 2 of the *OVOA1* gene. The *ovoa1-5* allele contains a premature stop codon and three *Eco*RV restriction sites in exon 2 of the *OVOA1* gene, accompanied by insertion of the homologous template for editing of the *ALS1* gene (highlighted in yellow) and duplications of the flanking genomic DNA (highlighted in blue). The *ovoa1-6* allele contains a premature stop codon TAG with two *Nhe*I restriction sites (GCTAGC, underlined) and one *Eco*RV restriction site in exon 3 of the *OVOA1* gene, accompanied by insertion of the *ALS1* homologous template and duplications of the flanking genomic DNA. Presumably, changes in the *ovoa1-4* allele were introduced by homologous recombination, while *ovoa1-5* and *ovoa1-6* alleles were edited by a mixed mechanism combining homologous recombination and non-homologous end-joining, with the latter typically accompanied by additional insertions, deletions, or duplications.

##### Editing of the selectable marker gene *ALS1* (Cre09.g386758_4532.1)

*als1-1* (mutation also present in the *ovoa1-5* and *ovoa1-6* mutants), exon 8

TGTGATCAAGGAGGCCTTTTACCTGGCCCGCACCGGCCGGCCCGGCCCTGTGCTGGTGGACGTGCCCACGGACATCCAGCAGCAGCTGGCGGTGCCGGACTGGGAGGCGCCCATGAGCATCACGG

*als1-2* (mutation also present in the *ovoa1-4* mutant), exon 8

TGTGATCAAGGAGGCCTTTTACCTGGCCCGCACCGGCCGGCCCGGCCCTGTGCTGGTGGACGTGCCCACGGATATCCAGCAGCAGCTGGCGGTGCCGGACTGGGAGGCGCCCATGAGCATCACGG

##### Editing of *OVOA1* (Cre08.g380000_4532.1)

*als1-2 ovoa1-4* (strain SSJL-1), exon 2

CTGTTTGAGACGGGCGTGGATGAGATGAGTTAAGATATCCTGTCGCGCGGGCGCGACGACTGGCCGCCCGTGCGCGAGGTGACCGAGTACCGCCGCAAGGCCTACGAGGTGGTGCGCGACGTCATCCTCAACCACCCCGCGCTCGACAAGCCCGAG

*als1-1 ovoa1-5* (strain SSJL-2), exon 2

CTGTTTGAGACGGGCGTGGATGAGATGAGTTAAGATATCCTGTCGCGCGGGCGCGACGACTGGCCGCCCCGGCCGGCCCGGCCCTGTGCTGGTGGACGTGCCCACGGATATCCAGCAGCAGCTGGCGGTGCCGGACTGGGAGGCGCCCCGCGGGCGCGACGACTGGCCGCCCGTGCGCGAGGTGGAGATGAGTTAAGATATCCTGTCGCGCGGGCGCGACGACTGGCCGCCCGTGCGCGAGGTGTTGGGACGACCTGTCGCGCGGGCGCGACGACTGGCCGCCCGTGCGCGAGGTGACCGAGTACCGCCGCAAGGCCTACGAGGTGGTGCGCGACGTCATCCTCAACCACCCCGCGCTCGACAAGCCCGAG

*als1-1 ovoa1-6* (strain SSJL-3), exon 3

ATCGGCTGGGACGATCCCGCCTTCCAGTCCGGCACCGCCAGCTGCTGCTGGATATCCGTGGGCACGTCCACCAGCACAGGGCCGGGCCGGCCGGTGCGGCCCCGCCTCGACCCAGATCGGCTGGGACGATCCCGCCTAAGCTAGCTTCCCCGCCTCGACCCAGATCGGCTGGGACGATCCCGCCTAAGCTAGCTTCATGGGCTTTGAGCACGAGCGCATCCACATCGAGACCAGCTCTGTGCTCATCCGCGAG

#### Supplementary Dataset S2: Sequence of His-tagged OVOA1 used for recombinant protein production.

The sequence shaded in grey contains a His_6_ tag used for purification. See Supplementary Figure S3 for more information.

MGSSHHHHHHSSGLVPRGSHMASANRVDTGACRNPSLNIRPDLARAANAITGAQVDDELVGPRGDWWWTGKRPEECPGFDKAAGVLRSLPLPNTRSFTRQSVLDYFDNSWTLTEVLFASLQTTDAFIRQPYHQLRHPMMFYYGHPAVLYINKFRVGGLLTDGLNQFFEQLFETGVDEMSWDDLSRGRDDWPPVREVTEYRRKAYEVVRDVILNHPALDKPEIGWDDPAFAVFMGFEHERIHIETSSVLIRELPLTSVRKPEFWPDYHPTSHNSSAPVPTQGVDFPVNNLVPVAGEKVVLGKDIAYPSFGWDNEYGAKEVNVQSFKATQFKVTNGEFLKFVKSSGYSNPKYWSAEGWGWKTFRNVKWPTFWMPDGPQGLHRYKLRVLFDAIDMRWDWPVDANYHEARAFAAWRTEADGASVHYRLITEAEHNLLRNSRDRVDAHLHTKAGSAQAAAAAAEPGRRAGANERVSPDLAMELSGANATARDAASGAGGFNVQLAHSSQNPVTELPPSEKGFYDTFGSAWEWAEDHYAAFPGFKVHPFYEDFSAPCFAGKHQLILGGSFISTGQLASKFARYQFRPHFFQHATFRLVVPDTDLSLYDAARYGPSNPITPFFETSCMDSAPPHVGDGPCCSKARRQAFTPTVLEEAASTKAKLEAAQVAYESDAILSQYLSLHYGPVDRVYPELANETGIIAAALDFPAKLAECLSSWAERAGVLPAAAADGASASGSGRALDLGCAVGRSTFELGRRFGEVVGVDISKTFIEVAAKIRDNGRMDYECAVEGETTERLTAALPAASAAAAGRCRFLQGDATSLPASSPTAPGSFDAVLAANLLDRVPEPATCLAQIKAALKPGGVALLTSPFSWLEQYTDRSNWLGGRYVDGLARRSADALKAAVAELGFEVLEEGSMPLIIREHGRKYQLINAHKLVMRLKA
